# Nuclear RNA export factor variant initiates piRNA-guided co-transcriptional silencing

**DOI:** 10.1101/605725

**Authors:** Kensaku Murano, Yuka W. Iwasaki, Hirotsugu Ishizu, Akane Mashiko, Aoi Shibuya, Shu Kondo, Shungo Adachi, Saori Suzuki, Kuniaki Saito, Tohru Natsume, Mikiko C. Siomi, Haruhiko Siomi

## Abstract

The PIWI-interacting RNA (piRNA) pathway preserves genomic integrity by repressing transposable elements (TEs) in animal germ cells. Among PIWI-clade proteins in *Drosophila*, Piwi transcriptionally silences its targets through interactions with cofactors, including Panoramix (Panx) and forms heterochromatin characterized by H3K9me3 and H1. Here, we identified Nxf2, a nuclear RNA export factor (NXF) variant, as a protein that forms complexes with Piwi, Panx, and p15. Panx-Nxf2-p15 complex formation is necessary in the silencing by stabilizing protein levels of Nxf2 and Panx. Notably, ectopic targeting of Nxf2 initiates co-transcriptional repression of the target reporter in a manner independent of H3K9me3 marks or H1. However, continuous silencing requires HP1a and H1. In addition, Nxf2 directly interacts with target TE transcripts in a Piwi-dependent manner. These findings suggest a model in which the Nxf2-Panx-p15 complex enforces the association of Piwi with target transcripts to trigger co-transcriptional repression, prior to heterochromatin formation in the nuclear piRNA pathway.

**Highlights:**

1. Nxf2 plays an essential role in the Piwi–piRNA pathway
2. Formation of Piwi-Panx-Nxf2-p15 (PPNP) complexes stabilizes both Panx and Nxf2
3. The PPNP complex triggers transcriptional silencing before heterochromatin formation
4. Nxf2 directly binds to target transcripts in a Piwi-dependent manner

## Introduction

Transposable elements (TEs) act as endogenous mobile mutagens to alter the sequence and structure of the genome, thereby changing the transcriptome and chromatin structure. TEs are often deleterious to the host by, for example, disrupting a gene, but may also be adaptive and drive genome evolution (Chuong et al., 2017; Han and Boeke, 2005). TEs comprise nearly half of the human genome and approximately 30% of the genome of *Drosophila melanogaster*; thus, the successively arising families of TEs are the main drivers of genome expansion. In metazoans, TEs are silenced by the piRNA pathway, in which Piwi proteins are guided by Piwi-interacting RNAs (piRNAs) to their targets (Ernst et al., 2017; Iwasaki et al., 2015; Ozata et al., 2019). The piRNA pathway is also essential for germline development and fertility in animals (Carmell et al., 2007; Cox et al., 1998; Houwing et al., 2007; Lin and Spradling, 1997).

In the *Drosophila* ovary, two cytoplasmic PIWI paralogs, AGO3 and Aubergine (Aub), engage an amplification loop termed the “ping-pong cycle” to cleave both TE transcripts and long piRNA precursor transcripts arising from piRNA clusters, which comprise a large number as well as various types of fragmented TEs, leading to the post-transcriptional silencing of TEs and production of piRNAs (Brennecke et al., 2007; Gunawardane et al., 2007). These piRNAs can in turn trigger the production of phased piRNAs from piRNA precursors, which generates a dazzling variety of piRNAs and is coupled to the activity of a third Piwi protein (Han et al., 2015; Homolka et al., 2015; Mohn et al., 2015). Phased piRNAs are also produced in ovarian somatic cells by a ping-pong cycle-independent mechanism, which are in turn also loaded onto Piwi (Han et al., 2015; Mohn et al., 2015). This loading initiates the transport of Piwi into the nucleus where it drives the transcriptional silencing of target TEs, by inducing specific histone modification and/or facilitating the folding of chromatin into a higher-order structure (Iwasaki et al., 2016).

Lack of the nuclear Piwi activity results in de-repression of TEs, which is concomitant with decreases in both H3K9me3 repressive epigenetic marks and RNA polymerase II (Pol II) occupancy on target TE loci (Donertas et al., 2013; Huang et al., 2013; Le Thomas et al., 2013; Ohtani et al., 2013; Rozhkov et al., 2013; Sienski et al., 2012). This suggests a scenario of how the nuclear piRNA pathway works: Piwi and its bound piRNAs scan for target TEs by complementary base-pairing with their nascent transcripts. Upon targeting, Piwi recruits chromatin factors including H3K9me3 methyltransferases such as Eggless (also called dSetDB1) to initiate heterochromatin formation. The H3K9me3 modification is then bound by HP1a to maintain and propagate epigenetic silencing. However, thus far, direct association of the Piwi–piRNA complex with target transcripts and/or H3K9me3 methyltransferases has not been demonstrated.

Furthermore, depletion of Maelstrom (Mael), a Piwi cofactor in the nuclear piRNA pathway, does not decrease H3K9me3 levels at Piwi-targeted TE loci, suggesting that the repressive histone mark *per se* is not the final silencing mark for transcriptional gene silencing mediated by Piwi–piRNA complexes (Sienski et al., 2012). Recently, we identified linker histone H1 as a component of a nuclear Piwi complex and found that depletion of H1 de-represses Piwi-targeted TEs and their surrounding genes without affecting the density of H3K9me3 marks and HP1a at target TE loci (Iwasaki et al., 2016). Instead, we demonstrated that the chromatin accessibility at Piwi-targeted TE loci is modulated by H1. These findings suggested a model in which Piwi recruits H1 and forces it to stay on target TE loci to induce chromatin compaction, thereby repressing target TE transcription, and that the nuclear piRNA pathway adopts collaborated actions of H1 and H3K9me3 mark to maintain silencing of the TE state by modulating the chromatin state.

In addition to Mael and H1, biochemical and genetic analyses have also identified a number of putative Piwi cofactors including DmGTSF1/Asterix (Arx) and Panoramix (Panx)/Silencio (Brower-Toland et al., 2007; Czech et al., 2013; Donertas et al., 2013; Le Thomas et al., 2013; Muerdter et al., 2013; Ohtani et al., 2013; Sienski et al., 2015; Sienski et al., 2012; Wang and Elgin, 2011; Yu et al., 2015). These findings together suggest that multiple pathways leading to Piwi-mediated TE silencing may exist, and raise the question of whether these pathways are independently initiated or have a common initial event.

To investigate this issue, we reanalyzed Panx, which interacts with Piwi and promotes the deposition of H3K9me3 marks on target TE chromatin by H3K9me3 histone methyltransferase, Eggless (Sienski et al., 2015; Yu et al., 2015). Here, we biochemically isolated Nxf2, nuclear RNA export factor (NXF) variant, as a component of Panx-associated complexes. Nxf2 further associates with p15 (also called Nxt1), a co-adaptor for nuclear RNA export (Fribourg et al., 2001; Herold et al., 2001; Kerkow et al., 2012). The NXF family comprises four members in *Drosophila*, among which Nxf1 is an essential mRNA nucleocytoplasmic export factor (Bjork and Wieslander, 2014; Izaurralde, 2002; Stutz and Izaurralde, 2003; Wickramasinghe and Laskey, 2015). However, other members (Nxf2-4) are gonad-specific and their functions are not known, although they share common domain structures with Nxf1 (Herold et al., 2001; Herold et al., 2000; Herold et al., 2003). Detailed analysis of Nxf2 function in the nuclear piRNA pathway revealed that the interactions between Panx, Nxf2, and p15 are necessary to maintain the protein stability of Nxf2 and Panx. Moreover, ectopic targeting of Nxf2 initiates co-transcriptional repression of the target reporter gene prior to heterochromatin formation, and H3K9me3 marks and H1 are required at later time points. Notably, the RNA binding domain of Nxf2 is essential for recruitment of the complex to target TEs. CLIP experiments demonstrated that both Piwi and Nxf2 directly interact with Piwi-targeted TE transcripts and that the association of Nxf2 with target TE transcripts is Piwi-dependent. These results suggest that Nxf2 enforces the association of Piwi-Panx-Nxf2-p15 (PPNP) complexes with the nascent transcripts of target TEs, and triggers co-transcriptional repression in the nuclear silencing pathway. Our results provide an unexpected connection between an NXF variant and small RNA-mediated co-transcriptional silencing.

## Results

### Nxf2 forms a complex with Piwi and Panx, and plays an essential role in the piRNA pathway

To reveal the precise mechanism by which Piwi–piRNA complexes transcriptionally silence TEs, we raised a monoclonal antibody that specifically recognises Panx (Figure S1A). Using this antibody, Panx-associated complexes were immunopurified from OSCs, a cultured ovarian somatic cell line, in which piRNA-guided silencing operates (Saito et al., 2009). Mass spectrometry analysis of the purified complexes (Figure 1A, Table S1) revealed that, in addition to Piwi, Nxf2 is present in the associated complexes. Nxf2 is a variant of the nuclear RNA export factor (NXF) family, which harbours similar domain structures (Herold et al., 2001; Herold et al., 2000; Herold et al., 2003) (see also Figure S1H). Among NXF variants, Nxf1 is highly conserved, ubiquitously expressed, and well known to be involved in the nuclear export of various mRNAs (Bjork and Wieslander, 2014; Izaurralde, 2002; Stutz and Izaurralde, 2003; Wickramasinghe and Laskey, 2015). However, Nxf2 and Nxf3 are almost exclusively expressed in the ovary, while Nxf4 is specifically expressed in the testis (Gramates et al., 2017). NXF variants other than Nxf1 are not involved in general mRNA export (Herold et al., 2001; Herold et al., 2003) and their functions remain to be revealed. By using a specific monoclonal antibody generated against Nxf2 (Figure S1B), we confirmed the interaction among Panx, Piwi, and Nxf2 in both OSCs and *Drosophila* ovary (Figures S1C and S1D). Immunoprecipitation followed by nuclear extraction also detected Piwi-Panx-Nxf2 complex (Figures 1B, S1E, and S1F). In addition, Panx and Nxf2 co-localised at the nucleus (Figure S1G), suggesting that the complex is formed within the nucleus. The interaction of Panx with NXF variants is restricted to Nxf2 (Figures S1H and S1I). Thus, we focused on the function of Nxf2 in the Piwi–piRNA pathway.

**Figure 1.**
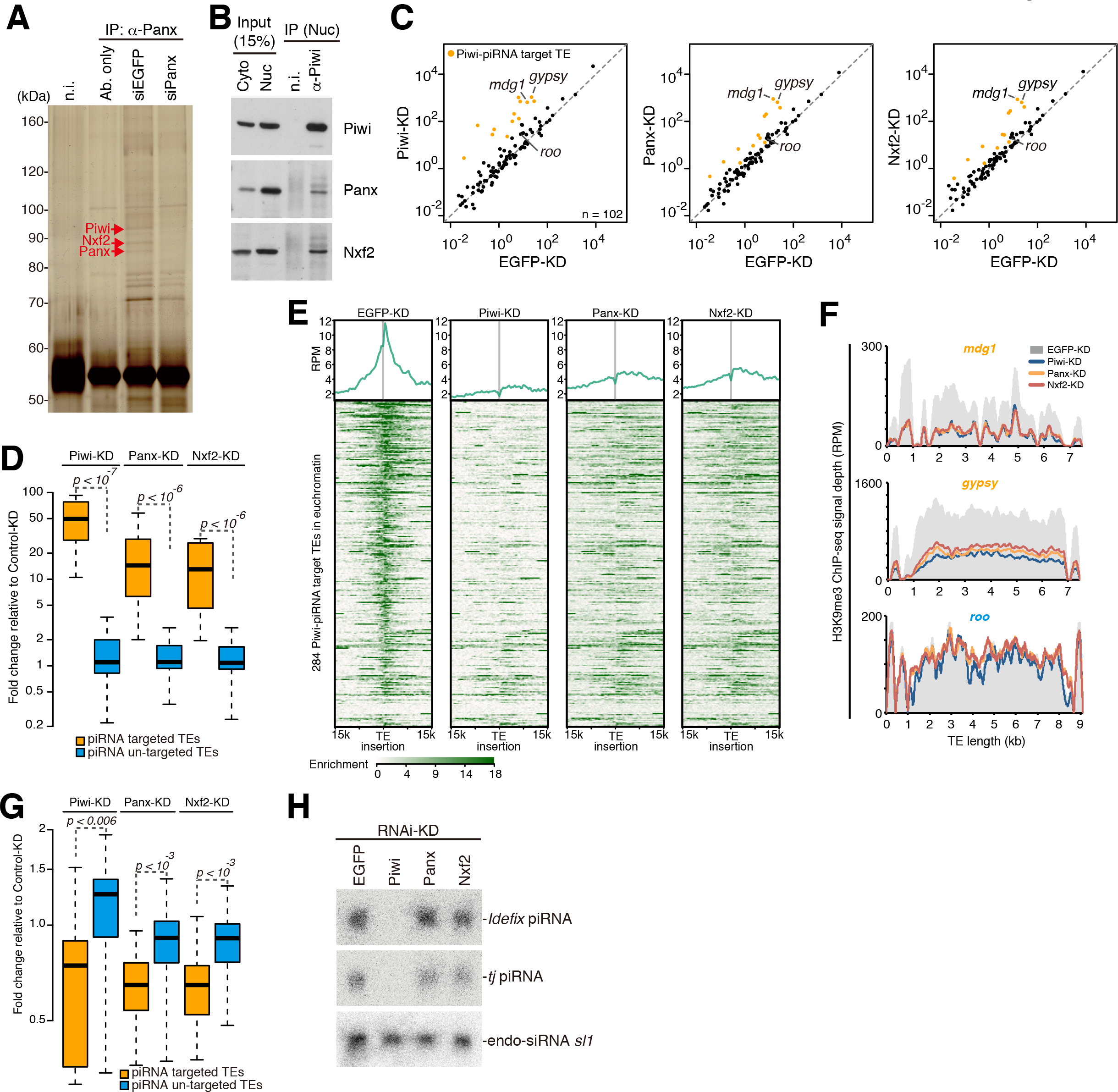
Nxf2 is required for Piwi–piRNA-mediated TE silencing in OSCs. **(A)** Immunoprecipitation (IP) from siRNA-transfected OSC lysate using anti-Panx antibody, followed by silver staining. Protein bands indicated by red arrows were identified by mass spectrometry analyses (see also Table S1). Mouse immunoglobulin G (IgG) (n.i.) was used for control IP. **(B)** IP from OSC lysate using anti-Piwi antibody, followed by Western blotting. Cyto: cytoplasmic fraction of OSCs. Nuc: nuclear fraction of OSCs. **(C)** Scatterplot of RPKM values for TEs in EGFP-(control), Panx-, or Nxf2-KD samples, based on RNA-seq. TEs for which the expression level differed from that of the control by more than tenfold in Piwi-KD (Piwi–piRNA-targeted TEs) are plotted in orange. Both x- and y-axis is a log10 scale. **(D)** Boxplots showing fold changes in the expression of Piwi–piRNA-targeted and un-targeted TEs based on RNA-seq upon the indicated KD. Piwi–piRNA-targeted TEs are as defined previously (Iwasaki et al., 2016). Boxplot whiskers show maxima and minima. *P*-values were calculated by Wilcoxon rank-sum test, and the y-axis is a log2 scale. **(E)** Meta-plot and heat map indicating H3K9me3 levels within 15 kb of the Piwi– piRNA-targeted TE insertion are shown for OSCs with the indicated KD. Heat map is sorted by the decrease of H3K9me3 levels in Piwi-KD against EGFP-KD. **(F)** Density plots for normalised H3K9me3 ChIP-seq signals over the consensus sequence from *mdg1*, *gypsy* (targeted by Piwi–piRNA, orange), and *roo* (not targeted by Piwi–piRNA, blue) TEs in EGFP-, Piwi-, Panx- and Nxf2-KD OSCs. **(G)** Boxplots as in **(D)** showing fold changes in the H3K9me3 levels of Piwi– piRNA-targeted and un-targeted TEs upon the indicated KD. **(H)** Northern blotting for *Idefix*-piRNA, *traffic jam* (*tj*)-piRNA, and esiRNA *sl-1* (control) on total RNA isolated from OSCs with the indicated KD.

To identify transcripts regulated by Nxf2, we performed mRNA-seq analysis of OSCs under Nxf2 knockdown (KD), and compared the results with those obtained for Piwi- and Panx-KD OSCs (Figures 1C, 1D, and S1J). TEs that were de-silenced upon Piwi depletion (Iwasaki et al., 2016) were also specifically de-silenced upon depletion of Panx or Nxf2. For example, *mdg1* and *gypsy* were among the most highly de-silenced TEs in Nxf2-, Panx-, and Piwi-KD experiments. Panx- and Nxf2-KD resulted in nearly identical patterns of TE de-silencing, albeit to a lesser extent than Piwi-KD, possibly because Piwi functions not only in TE silencing but also in piRNA biogenesis (see also Figure 1H). These results suggest that Nxf2 regulates TEs in the Piwi–piRNA pathway, in a manner resembling Panx, and are consistent with previous reports describing that Nxf2 was included in the list of candidate co-factors for the piRNA pathway identified by RNAi screening (Czech et al., 2013; Handler et al., 2013; Muerdter et al., 2013).

Depletion of Piwi has been shown to result in the reduction of H3K9me3 marks and H1 in the regions neighbouring TE insertions, resulting in de-repression of coding genes located near TEs (Iwasaki et al., 2016; Sienski et al., 2012). Coding genes that were de-repressed upon Nxf2-KD were highly similar to those observed in Piwi-KD, suggesting that Nxf2 also regulates genes located near TE insertions. The population of differentially expressed genes (DEGs) and their fold change were smaller in Nxf2-KD (Figures S1K–M), in line with the lower extent of TE de-silencing in Nxf2-KD (Figures 1C and 1D). These results indicate that Nxf2 functions in the Piwi–piRNA pathway, and does not affect the expression pattern of bulk mRNA, consistent with the previous finding that Nxf1 is the only member of the *Drosophila* NXF family that mediates the export of mRNAs (Herold et al., 2001; Herold et al., 2003).

We further analysed the H3K9me3 levels in these corresponding regions. ChIP-seq analysis of H3K9me3 marks at the euchromatic insertions of piRNA target TEs revealed that the depletion of Nxf2 decreased the H3K9me3 levels at these loci, similar to the case in Piwi or Panx depletion (Figures 1E–G). Notably, a severe decrease of H3K9me3 marks was observed not only at the TE insertions but also at the regions surrounding TE insertions (Figure 1E). The effect of Nxf2 or Panx depletion was weaker than that of Piwi depleation, and the results of Nxf2 and Panx depletion highly resembled each other, consistent with the effects on the expression levels of TEs observed by RNA-seq (Figures 1C and 1D). The analysis focusing on the TE consensus sequence confirmed that the effect on the H3K9me3 level was limited to the TEs targeted by Piwi–piRNA complexes (Figure 1G). Furthermore, the depletion of Nxf2 in OSCs did not affect the expression levels of piRNAs (Figure 1H), indicating that piRNA biogenesis including piRNA precursor export to the cytoplasm occurs in the absence of Nxf2. Together, these results indicate that Nxf2 is a key factor required for transcriptional silencing in the Piwi–piRNA pathway.

### Formation of Panx-Nxf2-p15 complex is essential for stabilization of Panx and Nxf2 protein levels

To further characterize the Piwi-Panx-Nxf2 complex, we immunopurified flag-tagged Nxf2 expressed in OSCs (Figure S2A), which was then subjected to shotgun proteome analysis (Figure S2B, Table S2). This analysis not only confirmed the presence of Panx and Piwi, but further detected p15, also known as Nxt1, in Nxf2-associated complexes (see also Figure S6C). We confirmed the interaction of Nxf2 with p15 in OSCs (Figure 2A), and p15 was also co-immunoprecipitated with Panx (Figure S2C). These results suggest the formation of a <u>P</u>iwi-<u>P</u>anx-<u>N</u>xf2-<u>p</u>15 complex, which we hereafter call the PPNP complex. p15 is a co-adaptor for nuclear RNA export and has been shown to form a heterodimeric complex with NXF proteins (Fribourg et al., 2001; Herold et al., 2001; Kerkow et al., 2012). Both Nxf1 and p15 are highly conserved from yeast to mammals, and the Nxf1–p15 complexes bind to mRNAs without strong sequence preference and have the ability to export mRNAs through the nuclear pore complex (NPC) (Herold et al., 2001; Levesque et al., 2001; Wiegand et al., 2002; Wilkie et al., 2001). Although a previous study showed that recombinant p15 can associate with Nxf1, Nxf2, and Nxf3 proteins, only the Nxf1–p15 complexes function in mRNA export, and the role of p15’s association with Nxf2 and Nxf3 remains unknown (Herold et al., 2001). We therefore analysed the involvement of p15 in Nxf2-mediated transcriptional silencing.

**Figure 2.**
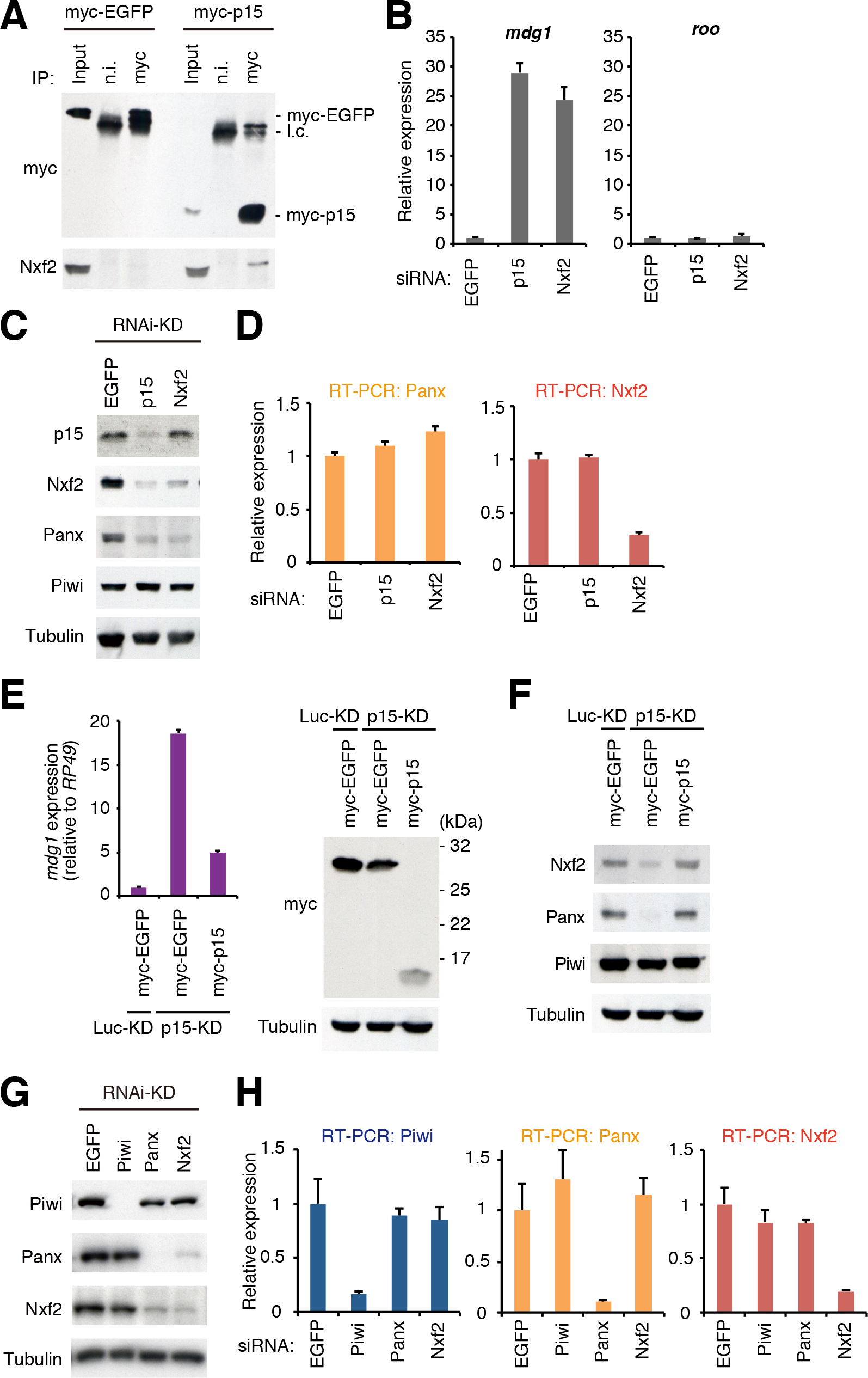
Panx-Nxf2-p15 complex formation is essential for the protein stability of Panx and Nxf2. **(A)** Western blot (WB) showing Nxf2 protein levels in the complex immunoprecipitated from OSCs expressing either myc–EGFP or myc–p15, using myc antibody. Mouse immunoglobulin G (IgG) (n.i.) was used for control immunoprecipitation. **(B)** RNA levels of *mdg1* and *roo* were quantified by qRT-PCR upon depletion of EGFP (control), p15, or Nxf2. Error bars indicate SD (n=3). Note that KD was performed for 2 days with a single siRNA transfection for this experiment. **(C)** WB showing p15, Nxf2, Panx, Piwi, and Tubulin (loading control) protein levels upon EGFP (control)-, p15-, or Nxf2-KD. **(D)** RNA levels of Panx and Nxf2 were quantified by qRT-PCR upon depletion of EGFP (control), p15, or Nxf2. Error bars indicate SD (n=3). **(E)** myc–EGFP or myc–p15 was expressed in Luc-(control) or p15-depleted OSCs, and *mdg1* expression levels were monitored by qRT-PCR (left panel). Expression values are normalised by the expression of *RP49*. Error bars indicate SD (n=3). Using the same samples, WB was performed using antibodies against myc and Tubulin (right panel). **(F)** myc–EGFP or myc–p15 was expressed in Luc-(control) or p15-depleted OSCs, and WB was performed using antibodies against Nxf2, Panx, Piwi, and Tubulin. **(G)** WB showing Piwi, Panx, Nxf2, and Tubulin (loading control) protein levels upon EGFP (control)-, Piwi-, Panx-, or Nxf2-KD. **(H)** RNA levels of Piwi, Panx, and Nxf2 were quantified by qRT-PCR upon depletion of EGFP (control), Piwi, Panx, or Nxf2. Error bars represent SD (n=3).

Depletion of p15 resulted in specific de-repression of Piwi–piRNA-targeted TE (Figure 2B). It was previously shown that p15 heterodimerises with human and *C. elegans* Nxf1 at its NTF2 (nuclear transport factor 2)-like domains (Fribourg et al., 2001; Klenov et al., 2014). We therefore analysed the interaction between p15 and Nxf2 mutant without NTF2-like domain (Figures S2D and S2E). This mutant could not associate with p15, and the rescue experiment showed that the NTF2-like domain of Nxf2 is indispensable for the silencing of *mdg1* TE (Figure S2F). These results suggest that the NTF2-like domain of Nxf2 associates with p15, as in the case of Nxf1, and this association is necessary for TE silencing.

Interestingly, depletion of p15 resulted in decreased levels of both Nxf2 and Panx proteins (Figure 2C). Since the RNA levels of Nxf2 and Panx remained unchanged in the p15-depleted samples (Figure 2D), p15 likely affects the stability of Nxf2 and Panx proteins. The protein level of p15 was unaffected by Nxf2-KD, possibly because p15 can bind to the other NXF proteins once Nxf2 is depleted (Figure 2C). De-silencing of the TEs and protein levels of Nxf2 and Panx could be rescued by expressing myc–p15 protein in p15-depleted OSCs (Figure 2E and 2F). Thus, p15 is an essential partner protein for Nxf2 to transcriptionally regulate Piwi–piRNA-targeted TEs, by stabilizing protein levels of Panx and Nxf2.

Notably, the knockdown of Nxf2 resulted in a severe decrease in the level of Panx protein (Figures 2C and 2G). A similar finding was observed in Panx-KD, where the level of Nxf2 protein was greatly decreased (Figure 2G). In contrast, neither Nxf2-nor Panx-KD affected the level of Piwi protein. RNA levels of Piwi, Panx, and Nxf2 in KD samples were determined by qRT-PCR, which showed that the levels of Panx and Nxf2 transcripts remained unchanged upon the knockdown of Nxf2 or Panx (Figure 2H). Destabilisation of Panx upon Nxf2-KD was also observed in the germ cells of the ovary (Figures S2G and S2H). Although the morphology of the ovary was not affected by the silencing of Nxf2 in the germline cells (Figures S2H), severe fertility defects were observed: flies with Nxf2 knockdown could lay eggs like the control flies, but none of them hatched (Figures S2I). This phenotype was similar to that observed in flies expressing Piwi only in ovarian somatic cells but not germ cells (Jin et al., 2013). Additionally, TEs targeted by piRNAs in the germline were specifically de-silenced in Nxf2-depleted flies (Figures S2J). These results together show that the Panx-Nxf2-p15 complex formation stabilizes the protein levels of Nxf2 and Panx (see also Figures 5 and S5), and this is essential for TE silencing.

### Tethering of Nxf2 to nascent mRNA leads to co-transcriptional silencing

We made use of reporter constructs to analyze the precise steps that the PPNP complex takes to mediate transcriptional silencing. First, we constructed reporter constructs whose transcripts are targeted by piRNAs derived from the *flam* locus, a prototype piRNA cluster (Brennecke et al., 2007; Post et al., 2014) (Figure S3A). The *luc* reporter gene containing *flam* fragments in sense orientation or in antisense orientation relative to the *luc* transcripts was transfected into OSCs; it was shown that only the antisense reporters were strongly silenced (Figure S3B-D). The antisense reporters were de-silenced in OSCs upon Piwi-KD (Figure S3E) or depletion of Piwi cofactors, including Panx and Nxf2 (Figure S3F), indicating that these factors were required for the co-transcriptional silencing in the Piwi–piRNA -targeted reporter system.

Previously, it was shown that the artificial recruitment of Panx can induce transcriptional silencing in *Drosophila* ovary (Sienski et al., 2015; Yu et al., 2015). For further functional analysis of Nxf2 in the Piwi–piRNA silencing pathway, we took advantage of the λN-boxB tethering system, in which λN fusion proteins are delivered to a reporter RNA *via* protein–RNA interaction (Baron-Benhamou et al., 2004). We established a tethering system using OSCs carrying a genome-integrated luciferase (*luc*) reporter driven by the ubiquitin promoter, which harbours 14 boxB sites in its intron (Figure 3A). Expression of λN-HA-tagged Nxf2 or Panx led to reporter silencing in a manner dependent on the λN peptide fused to Nxf2 or Panx (Figures 3B and S4A). Silencing induced by tethered Panx or Nxf2 results in strongly reduced *luc* RNA levels (Figure 3C). Knockdown of Panx and p15 weakened the repression by the enforced tethering of Nxf2, showing that Nxf2 needs Panx and p15 for efficient repression (Figure 3D). Consistent with previous reports (Sienski et al., 2015; Yu et al., 2015), the tethering of Piwi to the reporter failed to induce silencing (Figures 3B and 3C). λN-HA-myc-tagged Piwi localized at the nucleus (Figure S4B), indicating that λN-HA-myc-tagged Piwi harbored piRNA in OSCs (Saito et al., 2010; Yashiro et al., 2018). Because we were able to demonstrate that Piwi, which is guided to the target reporter *via* loaded piRNA, represses its targets (Figure S3), it is likely that only piRNA-guided Piwi is able to recruit silencing machinery to the target (see also Figure S7Q).

**Figure 3.**
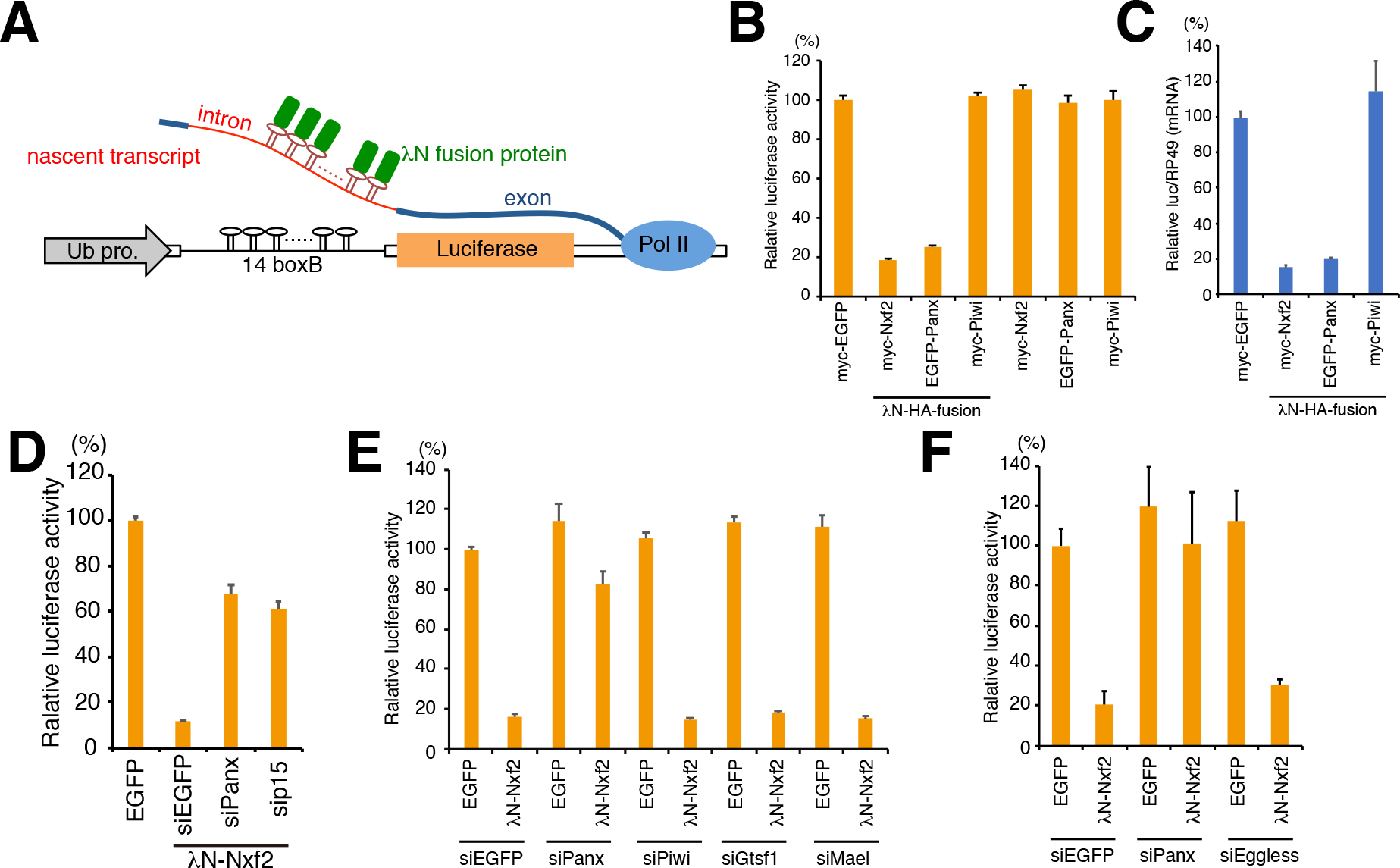
Nxf2 tethered to nascent mRNA causes co-transcriptional silencing. **(A)** Schematic of boxB-lN tethering system in OSCs. A genome-integrated luciferase reporter with 14 copies of boxB sites located within the intronic region of ubiquitin (Ub). λN fusion proteins are recruited to luciferase reporter RNA *via* 14× boxB. **(B)** Effects on luciferase activity of the proteins indicated below. OSCs carrying the reporter gene were transfected with plasmids expressing the indicated proteins and were harvested at 48 h post-transfection (hpt). Luciferase activities were normalised by the total protein amount. Bar graph shows luciferase activity relative to that of myc–EGFP (control). Error bars indicate SD (n=4). **(C)** qRT-PCR showing effects of the indicated λN fusion proteins on *Luc* mRNA level at 48 hpt. Levels of *Luc* mRNA were normalised by *RP49* mRNA. Bar graph shows the level of *Luc* mRNA relative to myc–EGFP (control). Error bars indicate SD (n=6). **(D)** Silencing by the enforced tethering of Nxf2 shows Panx and p15 dependent manner. Bar graph shows luciferase activity relative to that of the sample transfected with EGFP expression vector (control). Error bars indicate SD (n=3). **(E and F)** Effects of knockdown of the indicated genes on luciferase activity upon λN-Nxf2 expression at 48 hpt. Bar graph shows luciferase activity relative to that of the sample co-transfected with myc–EGFP and siEGFP (control). Error bars indicate SD (n=3).

To rule out the possibility that λN-Nxf2 interferes with the splicing of the reporter mRNA, we constructed a *luc* reporter with 10 boxB sites in the 3′ untranslated region (Figure S4C). Tethering of Nxf2 to the reporter mRNA also led to silencing in a λN peptide-dependent manner. To corroborate the results, we used an *in vivo* RNA-tethering system in the *Drosophila* ovary (Sienski et al., 2015). This assay builds on the α-*Tubulin* promoter driving the ubiquitous expression of a *GFP* reporter, which harbours five boxB sites in its 3′ untranslated region (Figure S4D). The expression of λN-Nxf2 with the MTD-Gal4 driver led to potent reporter silencing in germline cells but not in somatic cells of the ovary. Panx and Nxf2 proteins need to associate with each other to ensure their stability (Figure 2G and S2G); therefore, it is likely that tethering of Panx results in the recruitment of Nxf2, and *vice versa*. These results indicate that the Nxf2–Panx complex is sufficient to induce co-transcriptional silencing, when targeted to nascent RNA.

To dissect the mechanism of the observed co-transcriptional gene silencing in the tethering system in OSCs, we depleted the expression of genes known to be associated with Panx (Figure S1C) and involved in piRNA-directed transcriptional gene silencing. Depletion of Piwi or known Piwi cofactors [Gtsf1(Donertas et al., 2013; Ohtani et al., 2013; Sienski et al., 2012) and Mael (Sienski et al., 2012)] instead had no impact on the Nxf2-mediated repression (Figures 3E and S4E). Thus, Nxf2 acts with Panx most likely downstream from Piwi, Gtsf1, and Mael.

Furthermore, we examined the roles of chromatin-related factors in co-transcriptional silencing mediated by Piwi–piRNA complexes. Depletion of Eggless had no effect on the silencing by λN-Nxf2 in the tethering system using OSCs (Figures 3F and S4F). In addition to Eggless-KD, H1-KD and HP1-KD also had no impact on the silencing by Piwi–piRNA and λN-Nxf2 in the reporter system based on plasmids (Figures S3F and S4G-J). While this result is consistent with a report of a study using a plasmid-based reporter assay to detect Piwi-directed silencing in *Drosophila* ovary somatic sheet cells (Clark et al., 2017), it is at odds with previous reports demonstrating that the methyltransferase Eggless is required for Panx-mediated silencing using the tethering system in the *Drosophila* ovary (Sienski et al., 2015; Yu et al., 2015). These conflicting results are probably due to the different expression systems of λN fusion proteins. λN fusion proteins are driven by MTD-Gal4 constitutively in the tethering system in the ovary, whereas they are expressed transiently by DNA transfection in the tethering system using OSCs. The tethering system in the ovary probably enables us to observe late phases of a suppressed state of the reporter gene. In contrast, transient expression of λN fusion proteins makes it possible to analyse the mechanism of transition from initiation to the fully suppressed state of the reporter gene in our tethering system using OSCs. The results of Eggless-KD upon tethering of λN-Nxf2 suggest that the Nxf2– Panx pair elicits co-transcriptional repression prior to heterochromatin formation on TEs in the Piwi–piRNA pathway.

### Nxf2 triggers co-transcriptional repression prior to heterochromatin formation

To elucidate the precise mechanism by which λN-Nxf2 represses the reporter gene integrated in the genome, we examined the time course of silencing by λN-Nxf2, which was expressed in OSCs by transient transfection of expression vector. The expression level of λN-Nxf2 gradually decreased with a peak at 36 h post-transfection (hpt), and it had dropped below the detection limit at 96 hpt (Figure 4A). The luciferase activity reached its bottom at 48 hpt and remained suppressed at 96 hpt despite the absence of λN-Nxf2. Consistent with these observations, the level of Pol II association with the reporter gene decreased at 48 and 96 hpt (Figure 4B). Notably, no significant difference was observed in the occupancies of H1 and H3K9me3 at 48 hpt, despite λN-Nxf2 repressed the reporter gene (Figure 4A and 4C). This can be interpreted to mean that Nxf2-triggered co-transcriptional repression is independent of chromatin factors. By contrast, H1 and H3K9me3 accumulated on the reporter gene at 96 hpt (Figure 4C), suggesting that heterochromatin is formed at this time point on the reporter gene in the absence of λN-Nxf2, leading to continued silencing. To verify the role of heterochromatin formation in the gene silencing induced by λN-Nxf2, we knocked down H1 or HP1a after expression of λN-Nxf2. The effects of H1- and HP1a-KD on the λN-Nxf2-mediated silencing were negligible at 48 hpt (Figure 4D). To examine the roles of H1 and HP1a in the λN-Nxf2-mediated silencing at the latter time point, we determined luciferase activity at 96 hpt of the λN-Nxf2 expression vector followed by H1- and HP1a-KD. The depletion of H1 or HP1a partially restored luciferase activities upon tethering of λN-Nxf2 at 96 hpt (Figure 4E). These findings together suggest that λN-Nxf2 may initiate co-transcriptional repression prior to heterochromatin formation mediated by H1 and HP1a, which is required for maintaining the suppressed state of the reporter gene in the latter stage of silencing when little or no λN-Nxf2 is present. In other words, the silencing mode mediated by tethered Nxf2 may switch from Nxf2–Panx-dependent to heterochromatin-dependent.

**Figure 4.**
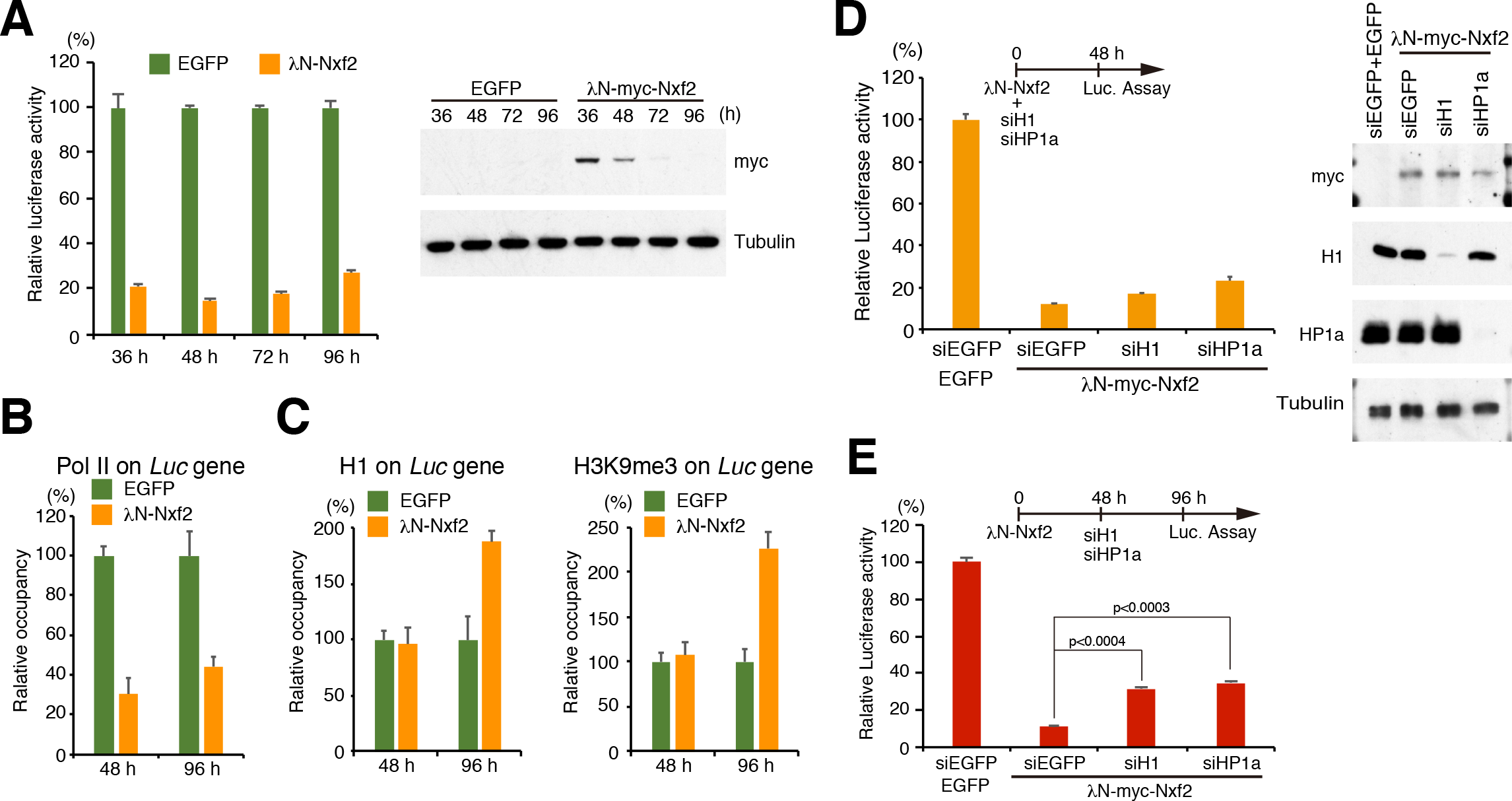
Nxf2 initiates co-transcriptional silencing before heterochromatin formation. **(A)** Time course of luciferase activity upon λN-Nxf2 tethering. Bar graph shows luciferase activity relative to that of EGFP at the indicated incubation time. Error bars indicate SD (n=3) (left panel). Expression levels of myc-tagged λN-Nxf2 were monitored using anti-myc antibody. Tubulin was used as a loading control (right panel). **(B)** ChIP-qPCR analysis of RNA polymerase II occupancy on the *Luc* gene upon λN-Nxf2 tethering. Bar graph shows the occupancy relative to that of the sample transfected with a plasmid expressing EGFP. Error bars indicate SD (n=3). **(C)** ChIP-qPCR analysis of histone H1 and H3K9me3 occupancy on the *Luc* gene. Bar graph shows the occupancy relative to that of the sample transfected with a plasmid expressing EGFP. Error bars indicate SD (n=3). **(D)** Effects of H1 and HP1a on luciferase activity upon λN-Nxf2 tethering at 48 hpt. OSCs were co-transfected with protein expression vectors and the indicated siRNAs and cultured for 48 h in accordance with the experimental scheme at the top of the figure. Bar graph shows luciferase activity relative to that of the sample expressing EGFP (control). Error bars indicate SD (n=3) (left panel). Western blots showing the protein levels of λN-myc–Nxf2, H1, HP1a, and Tubulin (loading control) in the same samples (right panel). **(E)** Effects of H1 and HP1a on luciferase activity upon λN-Nxf2 at 96 hpt. Transfection schedule of plasmids and siRNAs is shown at the top of the figure. siRNA transfection was carried out at 48h, since the depletion effect reaches maximum after 48 hpt. Bar graph shows luciferase activity relative to that of the sample expressing EGFP (control). Error bars indicate SD (n=3).

### The first LRR region of Nxf2 harbors RNA binding activity and is necessary for piRNA-mediated transcriptional silencing

To reveal the role of Nxf2 in the PPNP complex-mediated co-transcriptional silencing, we characterized the regions of Panx and Nxf2 proteins necessary for their function. This revealed that the middle region (300–399 aa) of Panx is necessary for its interaction with Nxf2 (Figures 5A and 5B) whereas its N-terminal region (1–169 aa) interacts with Piwi (Figure 5B). Rescue experiments and ectopic tethering experiments with the deletion mutants confirmed that both regions of Panx are indispensable for the transcriptional regulation (Figures S5A and S5B). We also produced a series of Nxf2 deletion mutants (Figure 5C) and found that the C-terminal UBA (ubiquitin-associated) domain of Nxf2 is necessary for interaction with Panx (Figure 5D). Although Nxf1 also harbours a UBA domain, we did not detect interaction between Nxf1 and Panx, indicating that Panx specifically recognises the UBA domain of Nxf2 (Figure S1I). These results suggest that the UBA domain of Nxf2 and middle region of Panx interact with each other, and N-terminal region of Panx is linked to Piwi.

**Figure 5.**
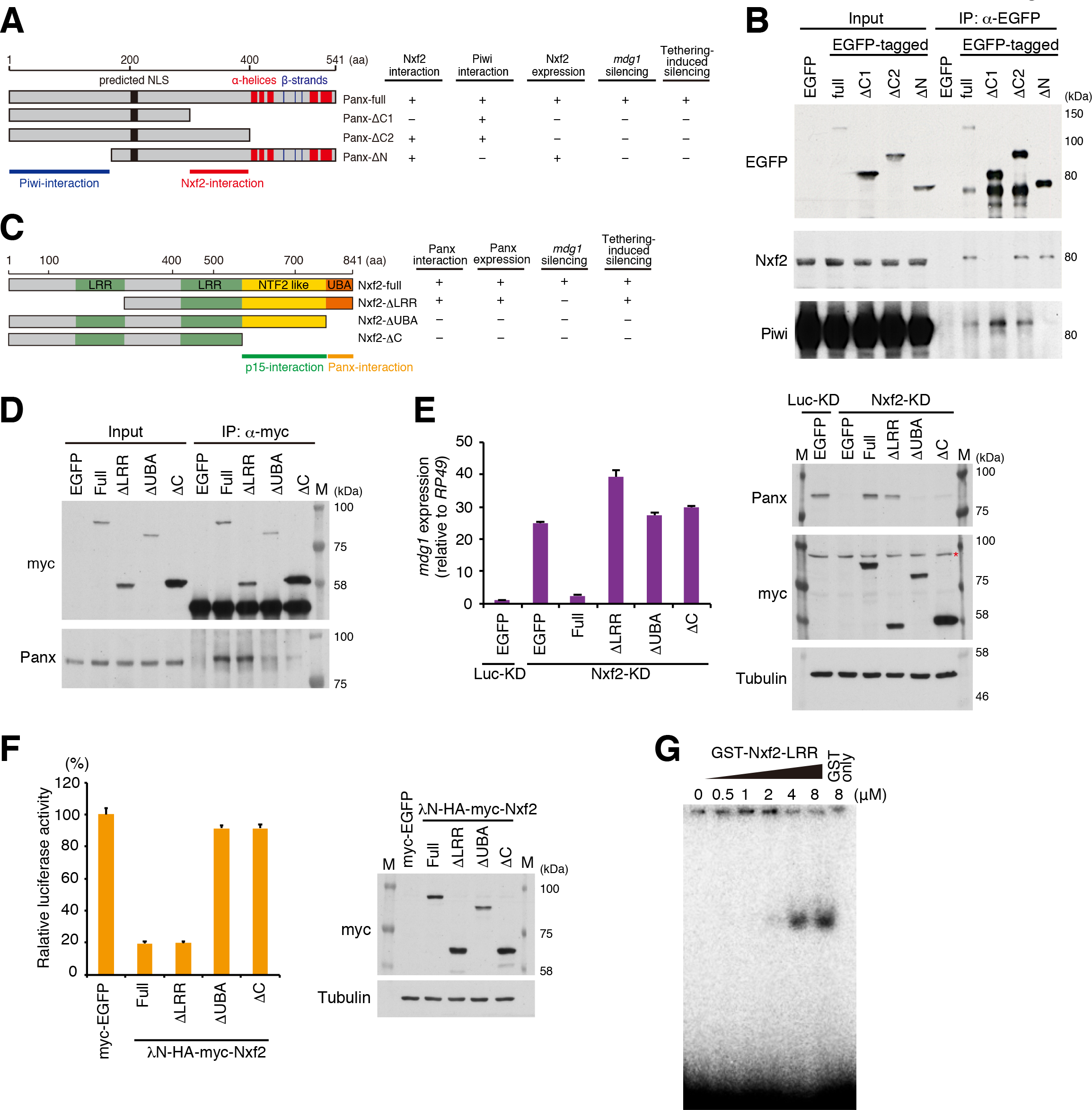
Nxf2 associates with Piwi–piRNA silencing complex to target RNA *via* its first LRR region. **(A)** Schematic of full-length Panx protein and deletion mutants. Predicted NLS and α-helices and β-strands are indicated (left panel). The results of rescue experiments are summarised (right panel). **(B)** Western blot (WB) of EGFP-Immunoprecipitation (IP) product from lysate of EGFP-tagged Panx deletion construct-expressing OSCs, using EGFP, Nxf2, and Piwi antibody. **(C)** Schematic of full-length Nxf2 and deletion mutant proteins. LRR: leucine-rich repeat domain, NTF2-like: nuclear transport factor 2-like domain, UBA: ubiquitin-associated domain (left panel). The results of rescue experiments and tethering assay are summarised (right panel). **(D)** IP using anti-myc antibody from lysate of OSCs expressing myc-tagged Nxf2 proteins, followed by WB using anti-myc or -Panx antibody. **(E)** *mdg1* expression levels were monitored by qRT-PCR in OSCs expressing exogenous Nxf2 proteins treated with either siLuc (control) or siNxf2 (left panel). Error bars indicate SD (n=3). Exogenous Nxf2 mRNAs are resistant to siRNA for Nxf2. Expression levels of Panx- and myc-tagged Nxf2 proteins were confirmed by WB using anti-myc and -Panx antibodies. Tubulin was used as a loading control. Asterisk indicates the background signal (right panel). **(F)** Effects of λN fusion Nxf2 proteins on luciferase activity at 48 hpt. Error bars indicate SD (n=3) (left panel). Expression levels of exogenous λN-HA–myc–Nxf2 proteins (right panel). **(G)** Electrophoretic mobility shift assay (EMSA) showing the binding of the first LRR region of Nxf2 (1–285 aa) to single-stranded RNA. The indicated amount of recombinant protein was mixed with 1 nM single-stranded RNA (16 nt).

Using the deletion mutants of Nxf2 (Figure 5C), we further performed rescue experiments by expressing each deletion mutant in Nxf2-depleted OSCs (Figure 5E). The expression of Panx protein could be stabilised not only by full-length Nxf2 but also by Nxf2-ΔLRR (leucine-rich repeat), showing that mutants that interact with Panx can rescue its protein level. However, Nxf2-ΔLRR, which associates with Panx and also Piwi (Figures 5D and S5C), could not recover the silencing of *mdg1* TE, suggesting that the LRR region of Nxf2 has an essential function other than formation of the PPNP complex. Consistent with this, piRNA-targeted reporters that were de-silenced upon depletion of Nxf2 (Figure S3F) could be rescued by full-length Nxf2, but not by Nxf2-ΔLRR (Figure S5D). We next investigated which of the Nxf2 mutants could induce silencing, when tethered to the reporter gene (Figure 5F). Interestingly, in addition to full-length Nxf2, Nxf2-ΔLRR mutant could induce silencing, indicating that the LRR region is dispensable once Nxf2 is tethered to the target gene. These results suggest that the LRR domain of Nxf2 is required to maintain recruitment of the PPNP complex to its targets.

It has been shown that Nxf1, the other NXF member in *Drosophila*, associates with bulk mRNA at its N-terminal RNA binding domain to induce its export (Viphakone et al., 2012). Additionally, a human homolog of Nxf2 associates with RNA through its N-terminal region (1–377 aa) (Herold et al., 2000). Meanwhile, both Nxf1 and Nxf2 in human use the N-terminal region to bind Alyref, a key factor for recruitment of Nxf1 to mRNA (Herold et al., 2000). To investigate the possible function of N-terminal region of *Drosophila* Nxf2 in piRNA pathway, we first preformed knockdown of Ref1 (Alyref homolog in *Drosophila*) in OSCs, but could not detect de-silencing of *mdg1* TE (Figure S5E). We further performed an electrophoretic mobility shift assay (EMSA) to analyse the RNA binding activity of the N-terminal LRR domain of Nxf2 and observed an association of single-stranded RNA with the purified GST-tagged first LRR domain of Nxf2 (1–285 aa) (Figure 5G). These results suggest that Nxf2 requires LRR domain, which harbours RNA binding activity, in addition to the other domains necessary for formation of the PPNP complex, in order to tether the silencing complex to its target transcripts.

### Nxf2 directly interacts with piRNA-target TE transcripts in a Piwi-dependent manner

The RNA-binding activity of Nxf2 prompted us to elucidate the mechanistic details of how Nxf2 is recruited to TE transcripts. We investigated cellular transcripts associated with Nxf2 and/or Piwi via UV crosslinking and immunoprecipitation (CLIP) assay (Flynn et al., 2015; Van Nostrand et al., 2016). We first raised a stable cell line of OSCs that express siRNA-resistant myc–Nxf2, since our Nxf2 antibody is not suitable for immunoprecipitation. The expression level of myc–Nxf2 is slightly higher than that of endogenous Nxf2, and Nxf2-siRNA specifically depletes endogenous Nxf2 (Figure S6A). myc–Nxf2 forms a complex identified using antibody against endogenous Panx (Figure S1C) and is capable of silencing *mdg1* TEs (Figures S6B and S6C). We therefore used this stable line to analyze the RNA population targeted by Nxf2.

We generated CLIP libraries using RNA isolated from myc–Nxf2 or Piwi immunoprecipitants. Immunoprecipitants were run on standard protein gels and transferred to nitrocellulose membranes, and a region 50 kDa above the protein size was excised for RNA isolation (Figure S6D), referring to the eCLIP method (Van Nostrand et al., 2016). The same region was excised for the input sample in order to generate a size-matched input (SMInput) library, which enables efficient background normalization and then leads to a more accurate measure of the enrichment of CLIP signals (Van Nostrand et al., 2016). Each CLIP experiment was performed in two biological replicates, and the reads were merged after checking the correlation of read counts of annotated peaks (Figure S6E). We first performed CLIP assay of myc–Nxf2 protein in UV-crosslinked and non-UV-crosslinked samples, to examine whether RNA associated with myc–Nxf2 could be captured by this assay. We observed a significant decrease in the amount of CLIP tags obtained for piRNA-targeted TE transcripts when CLIP was performed on non-UV-crosslinked samples (Figure S6F), indicating that we can specifically capture target RNA by the myc–Nxf2 CLIP assay.

Based on these results, we further performed myc–Nxf2 CLIP assay upon control (EGFP) or Piwi knockdown. Endogenous Nxf2 was also depleted, in order to increase the ratio of functioning myc–Nxf2. We first compared the enrichment of obtained CLIP tags to that of SMInput control, for the reads mapped in the sense or antisense direction of TEs (Figure 6A). This revealed that there was some fraction of TEs with enrichment of CLIP tags mapped in the sense direction, whereas this enrichment was not observed for the reads mapped in the antisense direction (Figure 6A). Additionally, we plotted the TEs targeted by Piwi–piRNA complexes, which showed that TEs with enriched CLIP tags were the targets of piRNAs (Figures 6A). Notably, the specific enrichment of myc– Nxf2 CLIP tags on piRNA-targeted TEs was lost upon the knockdown of Piwi (Figures 6B and S6G, see also Figure 6D), indicating that Piwi is required for Nxf2 to specifically associate with piRNA-targeted transcripts.

**Figure 6.**
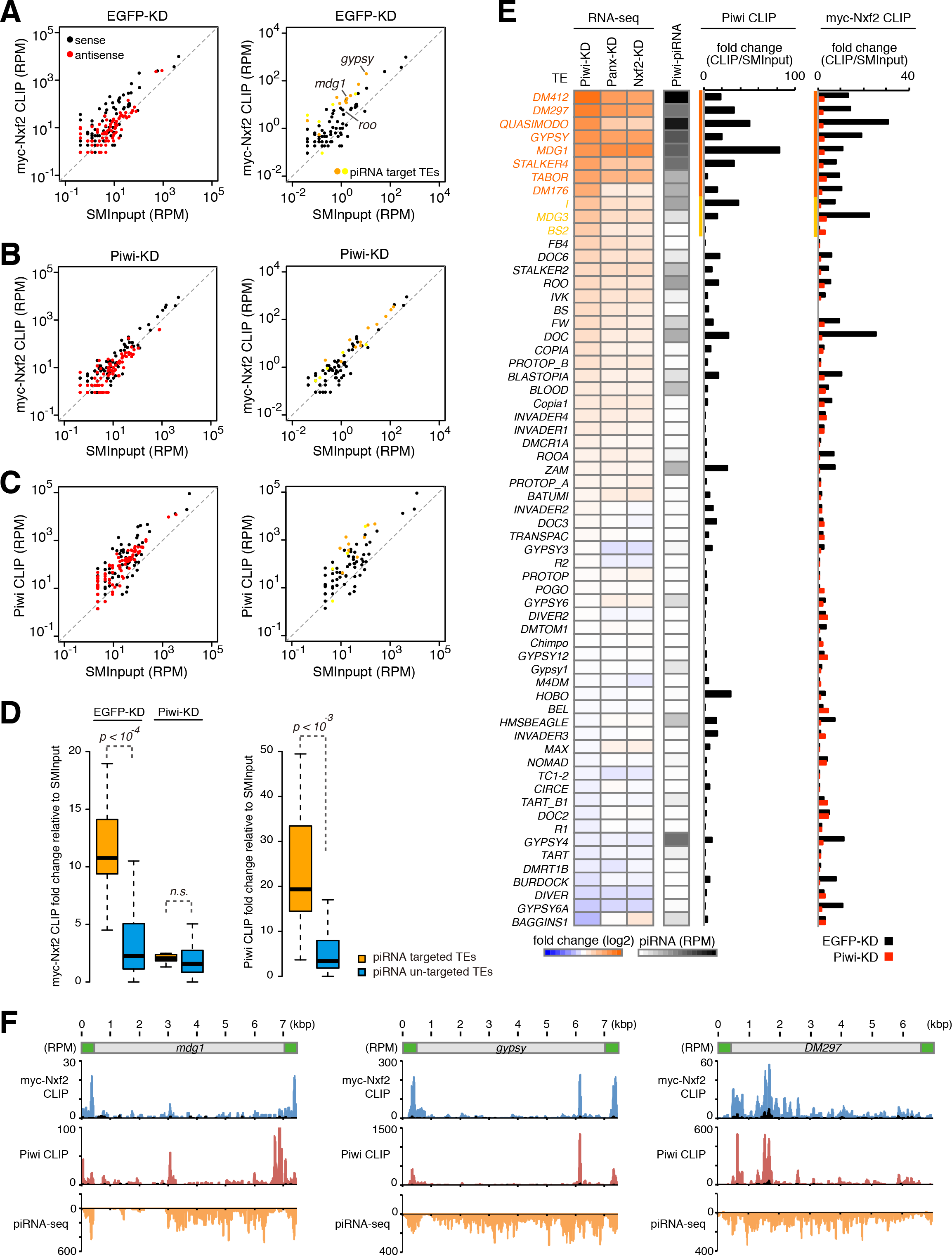
Nxf2 associates with nascent transcript of Piwi–piRNA-targeted TEs. **(A)** Scatterplot of RPM values for myc–Nxf2 CLIP tags (y-axis) and SMInput reads (x-axis) mapped to TEs in EGFP (control)-KD samples. Both x- and y-axes are log10 scales. Sense reads are plotted in black and antisense reads in red (left panel). TEs for which the expression level differed from that of the control by more than tenfold or fivefold in Piwi-KD (Piwi–piRNA-targeted TEs) are plotted in orange or yellow, respectively (right panel). **(B)** Scatterplot as in **(A)**. The plot was created using myc–Nxf2 CLIP data from Piwi-KD samples. **(C)** Scatterplot as in **(A)**. The plot was created using Piwi CLIP data. Note that RNAs used for creating Piwi CLIP libraries are in the size range of 43–73 nt, and therefore piRNAs are excluded. **(D)** Boxplots showing fold changes of the indicated CLIP tags to SMInput, for Piwi– piRNA-targeted and un-targeted TEs. Piwi–piRNA-targeted TEs are defined based on RNA-seq analysis (Figure 1). Reads mapped to sense direction of indicated TEs were used for calculation of CLIP tag fold change, and TEs with SMinput signal under 0.5 RPM were eliminated. Boxplot whiskers show maxima and minima. *p*-values were calculated by the Wilcoxon rank-sum test. **(E)** Heatmaps displaying fold changes of TE expression with indicated siRNA knockdown (normalized to EGFP-KD), and Piwi-associated piRNA levels obtained from small RNA-seq (RPM). The CLIP-seq diagram indicates fold changes of Piwi CLIP tags, and myc–Nxf2 CLIP tags relative to SMInput, obtained upon EGFP- or Piwi-KD. Reads mapped to sense direction of indicated TEs were used for calculation of CLIP tag fold change, and TEs with SMinput signal under 0.5 RPM were eliminated. Piwi-bound piRNA levels were determined by using reads mapped to antisense of the indicated TEs. **(F)** Density plots for myc–Nxf2 CLIP tags (blue plots), Piwi-CLIP tags (red plots), and Piwi–piRNA reads (orange plots) over the consensus sequence from *mdg1*, *gypsy*, and *DM297* (Piwi–piRNA target TEs). The LTR region of each TE is illustrated in green. SMInput is overlaid in black at each plot. y-axis has been adjusted to 100 for the plot of Piwi CLIP tags mapped to *mdg1*.

To further investigate the relationship between RNA targeting of Nxf2 and Piwi, we performed Piwi CLIP assay. As in the case of myc–Nxf2 CLIP, we observed the specific enrichment of CLIP tags against SMInput, specifically for those mapped in the sense direction of piRNA-targeted TEs (Figures 6C and 6D). We compared the expression level of TEs under Piwi-, Panx-, and Nxf2-KD, and also sequenced piRNA reads from Piwi-immunoprecipitant (Figure 6E). This revealed that TEs with enriched myc–Nxf2 and Piwi CLIP tags were de-silenced upon the depletion of Piwi, Panx, or Nxf2, and they were also targeted by piRNAs. A density plot of myc–Nxf2 and Piwi CLIP tags, along with piRNA reads, showed that myc–Nxf2 and Piwi CLIP tags had similar distributions, while piRNAs that can target wider regions of TEs were produced (Figure 6F). This indicates the possibility that specific piRNAs can silence certain transposons more efficiently than others. Since CLIP peaks tend to be detected on LTR regions or 5′ regions of transposons, piRNAs may silence their targets as soon as they are transcribed and therefore downstream RNAs may not be produced. These results, together with the finding that PPNP association is essential in order to stabilize the protein level of Nxf2 (Figure 2), suggest that the PPNP complex is recruited to nascent transcripts by base-pairing of Piwi-associated piRNAs with targets, and further enforced through the RNA binding activity of Nxf2.

## Discussion

Here, we propose a model in which the engagement of the PPNP complex with nascent TE transcripts initiates co-transcriptional silencing prior to heterochromatin formation (Figure S6H). Although the heterochromatin factors are not required for the initial step of the silencing (Figure 4), they play a significant role in maintenance of the repressed status in the later stages of TE silencing. Because the depletion of Piwi activity rapidly results in the de-repression of TEs (Figure 1), this also suggests that silencing initiation and heterochromatin formation undergo continuous cycling to enforce a silenced state of TEs. An important issue to be clarified next is the mechanism by which the PPNP complex initiates the silencing of TEs.

In *C. elegans*, NRDE-2 (nuclear RNAi defective-2) associates with the Argonaute protein NRDE-3 in the nucleus and is recruited by the NRDE-3/siRNA complex to nascent transcripts. This complex inhibits Pol II during the elongation, and directs the accumulation of Pol II at genomic loci targeted by RNAi (Guang et al., 2010). We investigated whether λN-Nxf2 directs the accumulation of Pol II at boxB sites in the co-transcriptional silencing (Figure S7A-C). Although the time course of Pol II occupancy upon λN-Nxf2 was examined by ChIP-qPCR at the boxB sites, we could not observe any accumulation of Pol II along the reporter gene.

Recently, it was reported that Pol II-associated proteins, PAF1 and RTF1, antagonize Piwi-directed silencing (Clark et al., 2017), which are factors involved in the modulation of promoter release, elongation, and termination of Pol II (Chen et al., 2015; Van Oss et al., 2017; Vos et al., 2018). The Panx-Nxf2-p15 complex might interfere with Pol II *via* the PAF1 complex, considering that fission yeast PAF1 represses AGO1/siRNA-directed silencing (Kowalik et al., 2015). Although we examined the effect of PAF1 and RTF1 on the co-transcriptional silencing, PAF1 and RTF1 do not appear to be involved in the co-transcriptional silencing mediated by λN-Nxf2 (Figures S7D-I). These results suggest that Pol II regulation by the PPNP complex differs from that of the small RNA-mediated Pol II regulation model proposed previously. It may be necessary to identify key factors involved in this regulation, possibly by performing proteome analysis of the unknown factors tethered to our reporter recruitment system.

We found a decrease in the active histone H3K4me3 marks on the reporter gene even at an earlier time point (48 hpt) at which the level of H3K9me3 marks remained low (Figure S7J and Figure 4C). Two reports about H3K4 methylation in Piwi-mediated silencing have been published. The level of H3K4 methylation on TEs is increased upon Piwi-KD (Klenov et al., 2014). The expression of artificial piRNAs that target a reporter locus induced transcriptional silencing associated with a decrease in the active H3K4-methylation marks (Le Thomas et al., 2013). In addition, the depletion of LSD1, which removes H3K4-methylation marks from promoters, had significant effects on the ability of Panx to repress the reporter gene (Yu et al., 2015). Therefore, we performed LSD1-KD and examined the effect of LSD1 on co-transcriptional silencing by the enforced tethering of Nxf2. However, the depletion of LSD1 had a limited impact on the silencing, whereas the level of *mdg1* was increased 27-fold upon LSD1-KD (Figures S7K-O). These findings together suggest that the decrease observed in H3K4me3 levels at 48 hpt in the tethering assay may not have been due to LSD1.

Nxf2 is a member of the NXF family, most members of which are known to associate with p15 (Herold et al., 2001). Although p15 interaction is essential for transcriptional silencing by Nxf2 (Figure 2), its molecular function remains to be revealed. In the case of Nxf1, its heterodimerization with p15 is essential for binding to NPC (Levesque et al., 2001; Wiegand et al., 2002). Considering that perinuclear chromatin positioning correlates with silenced expression and histone modifications (Harr et al., 2016), and that piRNAs are loaded onto Piwi at the Yb-body located near the nuclear membrane (Saito et al., 2010), it is interesting to analyze whether Piwi–piRNA silencing occurs near the nuclear periphery, and whether the association of p15 with Nxf2 would affect the localization of the Piwi–piRNA target chromatin in the nucleus. In addition, the NXF family is highly conserved in various species including humans (Bjork and Wieslander, 2014; Izaurralde, 2002; Stutz and Izaurralde, 2003; Wickramasinghe and Laskey, 2015). In the case of mouse, the expression of Nxf2 and Nxf3 is specific to the testis, instead of the ovary as in *Drosophila*, and male meiotic defects have been reported in mutant animals (Pan et al., 2009; Zhou et al., 2013). Taking these findings together with the fact that the mouse Piwi–piRNA pathway functions in testis rather than in ovary, it is possible that Nxf2 or other conserved NXF variants possess functions similar to *Drosophila* Nxf2 in mammals. In addition, Panx is not conserved in mammals (Sienski et al., 2015; Yu et al., 2015), so future studies of the proteins interacting with mouse Nxf2 may lead to the identification of a factor that functions in the place of Panx in mammals.

Our analysis revealed that PPNP complex plays a key role in co-transcriptional silencing in the Piwi–piRNA pathway by inducing multi-step transcriptional regulation (Figure S6H). Nxf2 is the core factor of this complex, and our results suggest that the nuclear export factor variant has been co-opted to repress TEs in the Piwi–piRNA pathway. We speculate that the co-option of Nxf2 into TE silencing is not coincidental but may reflect previous TE adaptation to exploit cellular mRNA transport pathways promoting the export of TE transcripts and replication/transposition (Checkley et al., 2013). Why is it necessary for Piwi to use the RNA binding activity of Nxf2 to silence their targets? Piwi, as a member of the Argonaute family, is thought to search for targets through random interactions between Piwi–piRNA and transcripts in the ocean of the transcriptome, many of which likely show partial complementarity with Piwi-loaded piRNAs (Klein et al., 2017). To distinguish “scanning Piwi” and “target-engaged Piwi,” the silencing pathway may need an engaged-in system to repress only targets and avoid unnecessary silencing of other cellular transcripts. It is possible that Nxf2 plays a role in supporting the association of Piwi that has found its *bona fide* targets (Figure S7P). Panx preferentially associates with Piwi, which loads piRNAs targeting TEs (Sienski et al., 2015), suggesting that there may be a system for Panx-Nxf2 to form the PPNP complex, specifically with Piwi that is engaged with its target TEs. Our CLIP analysis shows that Nxf2 cannot associate with target TEs without the activity of Piwi (Figure 6), indicating that the Panx-Nxf2-p15 complex itself does not recognize target transcripts, but rather recognizes target-engaged Piwi. In line with this, while piRNA-targeted Piwi can repress the reporter gene (Figure S3), λN-tethered Piwi cannot (Figure 3). This may be due to the Panx-Nxf2-p15 complex recognizing only piRNA-directed target-engaged Piwi, but not scanning or λN-tethered Piwi (Figures S7P and S7Q). Further studies will be necessary to elucidate the precise mechanism by which the Panx-Nxf2-p15 complex selectively recognizes only the “target-engaged Piwi” and forms a PPNP complex to induce silencing of its targets.

## Acknowledgements

We thank Wataru Ito for experimental support and critical discussions. We also thank Julius Brennecke for sharing plasmids; Kazumichi Nishida, Ryo Ohnishi, and Tatsuki Kinoshita for sharing protocols; and Soichiro Yamanaka for critical reading of the paper. This work was supported by funding from MEXT (17K08644) and Kato Memorial Bioscience Foundation to K.M.; from MEXT (15H05583, 17H05603, 18H02421, and 19H05268), Senri Life Science Foundation, NOVARTIS Foundation (Japan) for the Promotion of Science, and the Naito Foundation to Y.W.I.; from MEXT (17H05610) to S.A.; from Takeda Science Foundation to T.N.; and from MEXT (25221003) to H.S.

## Author Contributions

K.M. and Y.W.I. designed and performed most of the experiments with H.I., A.M., and A.S. S.K. and K.S. generated mutant flies for tethering assays, S.A. and T.N. performed LC-MS/MS analysis, and S.S. and M.C.S. generated Eggless antibody. Y.W.I. and H.S. conceived the study, and K.M., Y.W.I., and H.S. wrote the paper with input from the other authors.

