## Supplemental Information for "Nuclear RNA export factor variant initiates piRNA-guided co-transcriptional silencing"

Figure S1

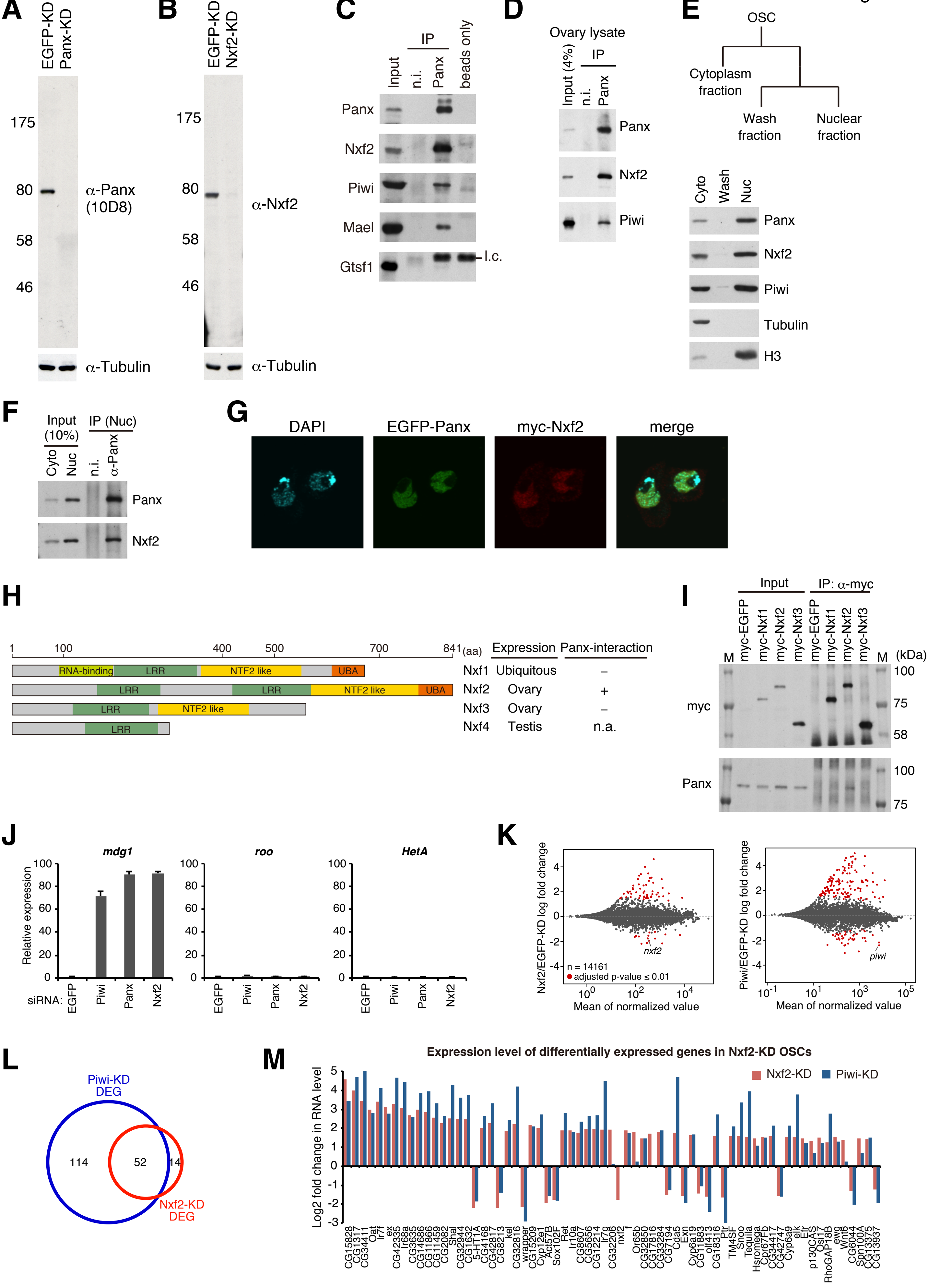
A
EGFP-KD
Panx-KD
175
80
58
46
α-Panx (10D8)
α-Tubulin
B
EGFP-KD
Nxf2-KD
175
80
58
46
α-Nxf2
α-Tubulin
C
Input
IP
n.i.
Panx
beads only
Panx
Nxf2
Piwi
Mael
Gtsf1
l.c.
D
Ovary lysate
Input (4%)
IP
n.i.
Panx
Panx
Nxf2
Piwi
E
OSC
Cytoplasm fraction
Wash fraction
Nuclear fraction
Cyto
Wash
Nuc
Panx
Nxf2
Piwi
Tubulin
H3
F
Input (10%)
IP (Nuc)
Cyto
Nuc
n.i.
α-Panx
Panx
Nxf2
G
DAPI
EGFP-Panx
myc-Nxf2
merge
H
1
100
400
500
700
841 (aa)
RNA-binding
LRR
NTF2 like
UBA
Expression
Panx-interaction
Nxf1 Ubiquitous –
Nxf2 Ovary +
Nxf3 Ovary –
Nxf4 Testis n.a.
I
Input
IP: α-myc
M
myc-EGFP
myc-Nxf1
myc-Nxf2
myc-Nxf3
(kDa)
myc
Panx
100
75
58
J
mdg1
roo
HetA
Relative expression
siRNA:
EGFP
Piwi
Panx
Nxf2
K
Nxf2/EGFP-KD log fold change
Piwi/EGFP-KD log fold change
nxf2
piwi
n = 14161
adjusted p-value ≤ 0.01
Mean of normalized value
L
Piwi-KD DEG
Nxf2-KD DEG
114
52
14
M
Expression level of differentially expressed genes in Nxf2-KD OSCs
Nxf2-KD
Piwi-KD
Log2 fold change in RNA level

**Figure S1. Related to Figure 1**.

**(A)** Western blotting (WB) shows the specificity of anti-Panx monoclonal antibodies raised in this study. Tubulin was used as a loading control.

**(B)** WB shows the specificity of anti-Nxf2 monoclonal antibodies raised in this study.

**(C)** Immunoprecipitation (IP) from OSC lysate using anti-Panx antibody, followed by WB of Panx, Nxf2, Piwi, Mael, and Gtsf1. IP-WB was performed under the same conditions as the silver staining described in Figure 1A. Mouse immunoglobulin G (IgG) (n.i.) was used for control IP. l.c.: light chain from the antibody.

**(D)** IP from fly ovary lysate using anti-Panx antibody, followed by WB of Panx, Nxf2, and Piwi.

**(E)** Scheme of preparation of cytoplasmic and nuclear fractions of OSCs. WB using Panx, Nxf2, Piwi, Tubulin, and histone H3 antibodies show the level of each protein in the separate fractions. Tubulin was detected in the cytoplasmic fraction, where histone H3 was enriched in the nuclear fraction.

**(F)** IP from nuclear OSC lysate using anti-Panx antibody, followed by WB.

**(G)** Immunofluorescence of OSCs co-transfected with EGFP-Panx- and myc-Nxf2-expressing vectors, using myc antibody (red). EGFP is detected as Panx signal, and DAPI staining (blue) shows the location of nuclei. EGFP-Panx and myc-Nxf2 co-localise in the nucleus.

**(H)** Schematic of NXF variants. LRR: leucine-rich repeat, NTF2-like: nuclear transport factor 2-like domain, UBA: ubiquitin-associated domain. Nxf1 is ubiquitously expressed, while Nxf2 and Nxf3 are almost exclusively expressed in ovary, and Nxf4 is specifically expressed in testis. The interaction between Nxf4 and Panx was not analysed, since Nxf4 expression is limited to testis, indicated as n.a. (not available).

**(I)** IP from lysate of OSCs expressing myc-tagged NXF variants, followed by WB using anti-myc and -Panx antibodies. M indicates protein markers. The results are summarised in the right panel in (H). Among NXF variants, only Nxf2 can interact with Panx.

**(J)** RNA levels of *mdg1*, *roo*, and *HetA* were quantified by qRT-PCR upon depletion of EGFP (control), Piwi, Panx, or Nxf2. Expression levels are normalised by the expression of *RP49.* Error bars represent SD (n=3). Piwi–piRNA-targeted TE was specifically de-silenced by Piwi-, Panx-, and Nxf2-KD, confirming the results of RNA-seq.

**(K)** MA plot of RPKM values (log10 scale) for mRNAs in the indicated KD samples, based on RNA-seq. Differentially expressed genes (DEGs) are in red.

**(L)** Venn diagram displaying the number of DEGs upon depletion of Nxf2 (red) or Piwi (blue).

**(M)** Log2 fold changes in mRNA levels of 67 DEGs in Nxf2-KD OSCs. mRNA levels calculated from RNA-seq data are used for Nxf2- and Piwi-KD OSCs. Most mRNAs whose expression was altered upon Nxf2-KD were also affected by Piwi-KD, suggesting that Nxf2 does not globally affect mRNA levels.

Figure S2

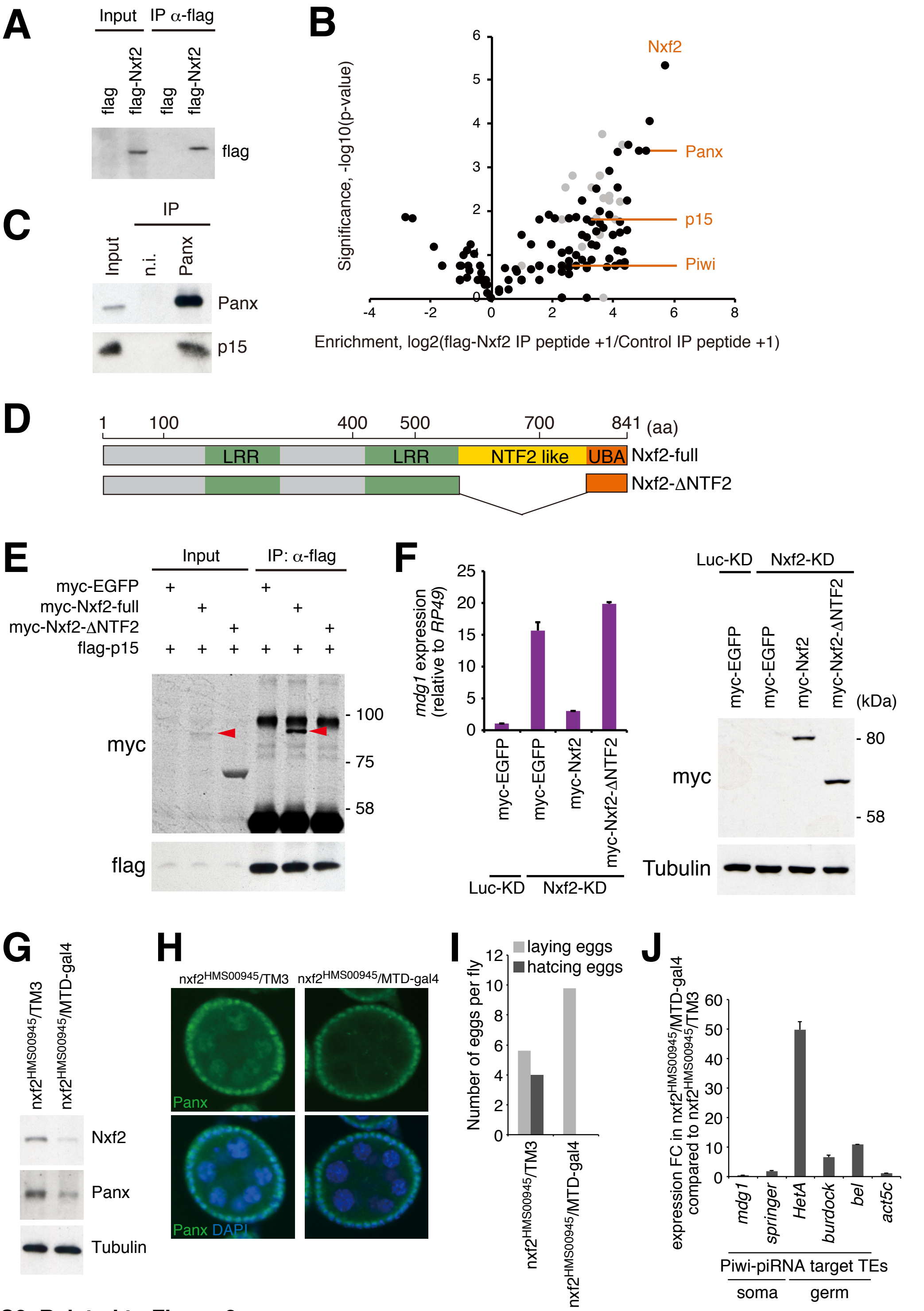

**Figure S2. Related to Figure 2.**

**(A)** Western blot (WB) showing flag–Nxf2 immunoprecipitant from OSCs expressing either flag only (control) or flag–Nxf2, using flag antibody. The immunoprecipitant was subjected to proteome analysis.

**(B)** Summary of the proteome analysis of peptide-eluted flag–Nxf2 component. The plot shows the enrichment of peptide obtained by flag-Nxf2 Immunoprecipitation (IP) over Control IP (log2 value of peptide count+1) on the x-axis, and significance calculated by replicate experiments as the p-value (-log10) on the y-axis. Enrichment and significance of each protein are listed in Table S2. Ribosomal proteins are plotted in gray.

**(C)** IP from OSC lysate using anti-Panx antibody, followed by WB of Panx and p15. IP was performed under the same conditions as silver staining described in Figure 1A.

**(D)** Schematic of full-length Nxf2 protein and deletion mutant of NTF2-like domain (Nxf2-ΔNTF2).

**(E)** Flag-IP performed using lysate of flag-p15- and myc-EGFP- or myc-Nxf2-ΔNTF2-expressing OSCs, followed by WB using myc or flag antibody. myc-Nxf2-full is indicated by a red arrowhead. p15 cannot interact with the Nxf2 construct lacking NTF2-like domain.

**(F)** myc-EGFP, myc-Nxf2, and myc-Nxf2-ΔNTF2 were expressed in Luc- (control) or Nxf2-depleted OSCs. *mdg1* expression levels were monitored by qRT-PCR. Expression values are normalised by the expression of *RP49*. Error bars indicate SD (n=3) (left panel). WB using an antibody against myc and tubulin using lysates from transfected OSCs (right). The ΔNTF2 deletion mutant, which cannot interact with p15, could not rescue the silencing of TE.

*(legend continued on next page)*

**Figure S2. Related to Figure 2. (continued)**

**(G)** WB of ovaries from nxf2HMS00945/MTD-gal4 flies using Nxf2, Panx, and tubulin (control) antibody. Note that the expression of shRNA is limited to germline cells by MTD-gal4. Depletion of Nxf2 in the germline cells also results in a decrease of Panx protein level.

**(H)** Immunofluorescence of ovaries from nxf2HMS00945 flies using Panx antibody (green). DAPI staining (blue) shows the location of nuclei. Decrease of Panx in Nxf2-depleted ovaries was observed specifically in the germ cells.

**(I)** Numbers of eggs laid and hatched per fly are indicated. Nxf2 depletion in Drosophila ovaries results in sterility phenotype.

**(J)** mRNA levels of Piwi–piRNA-targeted TEs and *actin 5c* (control) were quantified by qRT-PCR. Expression values are normalised by the expression of *RP49*. Error bars represent SD (n=3). Piwi–piRNA-targeted TEs in the germ cells were specifically de-silenced upon germline-specific Nxf2 depletion.

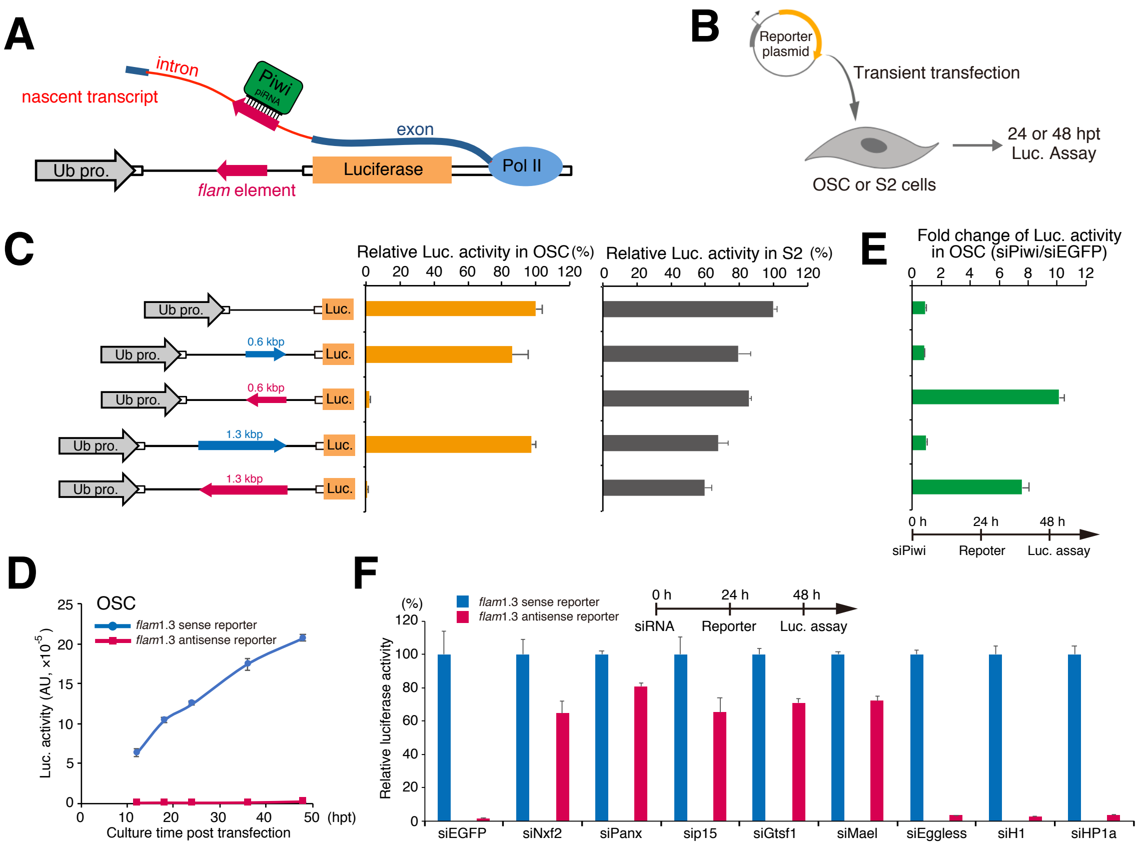

#### Figure S3. Related to Figure 3.

**(A)** Schematic of piRNA reporter gene assay. Reporter plasmid carries *flam* element in its intron. *Luc* mRNA harboring *flam* element in the antisense direction is targeted by Piwi carrying piRNAs derived from the *flam* locus.

**(B)** Experimental scheme. OSCs or S2 cells were transfected with reporter plasmids transiently, and harvested at 24 or 48 hpt.

**(C)** Left panel shows reporter gene constructs harboring *flam* elements (0.6 kbp or 1.3 kbp) inserted in the intron. Blue and red arrows indicate sense and antisense directions, respectively. Middle graph shows luciferase activity relative to that of reporter gene without *flam* elements in OSC. Right graph shows luciferase activity relative to that of reporter gene without *flam* elements in S2 cells. Error bar indicates SD (n=4).

**(D)** Time course of luciferase activity in OSCs. Blue and red arrows indicate reporter gene bearing a flam element (1.3 kbp) in the sense or antisense direction, respectively. Error bar indicates SD (n=3).

**(E)** Silencing of reporter gene harboring *flam* element (1.3 kbp, antisense direction) occurs in a Piwi-dependent manner. Graph shows fold change of luciferase activity in OSCs treated with siPiwi or siEGFP (control). Error bar indicates SD (n=3).

**(F)** Effects of knockdown of the indicated genes on piRNA reporter activity (1.3 kbp, *flam*). Red bar shows the luciferase activities of the antisense reporter relative to that of the sense reporter (blue). Error bars indicate SD (n=3). Transfection schedule of siRNA and reporter plasmids is shown at the top of the figure.

**Text related to Figure S3:** To elucidate the function of Nxf2 in the co-transcriptional silencing, we introduced another reporter construct, whose transcripts are targeted by Piwi-piRNA complexes derived from the *flam* locus, a prototype piRNA cluster (Brennecke et al., 2007) (Figure S3A). We selected 0.6 kbp and 1.3 kbp *flam* elements, from which a number of piRNAs are derived (Post et al., 2014), and cloned them into the intron of the *luc* reporter gene in the sense orientation or the antisense orientation relative to the *luc* transcripts. The sense reporters were expressed similar to control reporter gene lacking the *flam* element, whereas the antisense (piRNA-targeted) reporters were completely silenced, even though we introduced the reporter gene as a plasmid into OSCs (Figures S3B and S3C). The piRNA-targeted reporters were silenced immediately after transfection and continue to be suppressed (Figure S3D). In contrast, all reporters were expressed at a similar level to the control reporter gene in S2 cells (Figure S3C). The piRNA-targeted reporters were de-silenced in OSCs upon Piwi-KD (Figure S3E). These results suggest that Piwi guided to the target *via* its loaded piRNAs can recruit silencing machinery to the target, but not Piwi tethered by λN-peptide to boxB (Figure 3B and 3C). Then, we depleted the expression of Nxf2, Panx, p15, and known cofactors Gtsf1 and Mael, which de-silenced the piRNA-targeted reporter in OSCs (Figure S3F), indicating that these factors were required for the co-transcriptional silencing in the Piwi-piRNA-targeted reporter system. Notably, the reporter genes do not have to be integrated into the genome, suggesting that the co-transcriptional silencing occurs independent of the chromatin context. Furthermore, we examined the roles of chromatin-related factors on co-transcriptional silencing mediated by Piwi-piRNA complexes. Eggless-KD, H1-KD, and HP1-KD also had no impact on the silencing by Piwi-piRNA in the reporter system based on plasmids (Figure S3F).

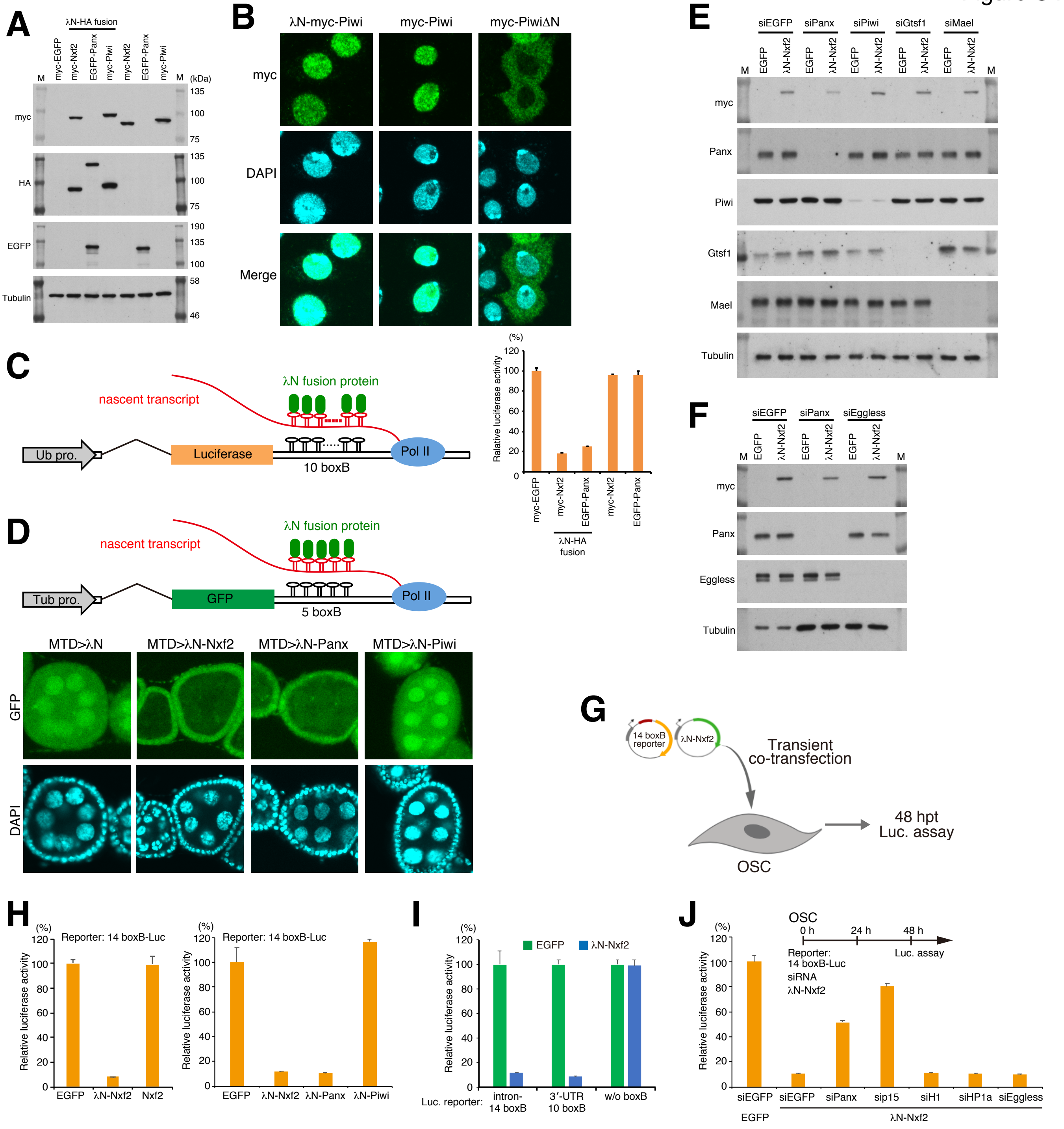

#### Figure S4. Related to Figure 3.

**(A)** Western blotting (WB) associated with Figure 3B shows the exogenous expression of λN-fused proteins. Nxf2 and Piwi were tagged with λN-HA and myc, whereas Panx was tagged with λN-HA and EGFP, since the expression level of EGFP–Panx was higher than the level of myc-tagged Panx, which is quite unstable. M indicates protein markers.

**(B)** Immunofluorescence of OSCs transfected with λN-myc-Piwi-, myc-Piwi-, or myc-PiwiΔN-expressing vectors, using myc antibody (Green). DAPI staining (blue) shows the location of nuclei. myc-PiwiΔN lacking nuclear localization signal (NLS) was distributed in cytoplasm (Saito et al., 2010; Yashiro et al., 2018). λN-myc-Piwi localized in the nucleus similarly to myc-Piwi, indicating that λN-myc-Piwi harbored piRNA in OSCs, even though it failed to induce silencing of the boxB reporter gene (Figures 3B, see also Figure S4D, and S4H). This may be due to only Piwi, which is guided to the target *via* its loaded piRNAs, potentially being able to recruit silencing machinery to the target (see also Figure S3C and S7Q).

**(C)** Schematic of boxB-λN tethering system in OSCs shows a genome-integrated luciferase reporter with 10 copies of boxB sites located within the 3′ UTR of *luc* mRNA (left). Bar graph shows relative luciferase activity normalized by total protein amount at 48 h post-transcription (right). Error bars indicate SD (n=4). Tethering of Nxf2 and Panx to the 3′ UTR of *luc* mRNA leads to silencing in OSCs.

*(legend continued on next page)*

#### Figure S4. Related to Figure 3. (continued)

**(D)** Schematic of boxB-λN tethering system in fly ovary shows a genome-integrated *GFP* reporter with five copies of boxB sites located within the 3′ UTR of *GFP* mRNA. λN-fused proteins are driven by MTD-Gal4. Tub pro: *α-Tubulin* gene promoter (top panel). Confocal images depict GFP fluorescence and DAPI signals in egg chambers expressing the indicated λN-fusion proteins in the germline. Expression of λN-Nxf2 and Panx leads to reporter silencing in germ cells, but that of λN-Piwi does not. This is consistent with our data in Figure 3B and previous reports (Sienski et al., 2015; Yu et al., 2015).

**(E and F)** Results of WB associated with Figures 3E and 3F, showing the expression of λN-HA–myc–Nxf2 and the effect of siRNAs on target gene expression.

**(G)** Experimental design. According to Figure S3, the reporter genes do not have to be integrated into the genome, suggesting that the co-transcriptional silencing occurs independent of the chromatin context. To examine this issue, we carried out co-transfection of reporter plasmid (14 boxB-Luc) and expression plasmids for λN-fusion protein to OSCs, transiently. Cells were harvested at 48 hpt.

**(H)** Effect on luciferase activity of the proteins indicated below. Error bars indicate SD (n=4). Even if a plasmid with 14 boxB reporter sites was transiently introduced into OSCs, λN-Nxf2 repressed luciferase activity.

**(I)** λN-Nxf2 represses luciferase activity of reporter plasmid harboring 10 boxB sites in its 3′ UTR, but not that of reporter plasmid without boxB. Error bars indicate SD (n=4).

**(J)** Effects of knockdown of the indicated genes on boxB reporter activity upon λN-Nxf2 expression. Bar graph shows luciferase activity relative to that of the sample co-transfected with myc-EGFP and siEGFP (control). Error bars indicate SD (n=4). Transfection schedule of siRNA and reporter plasmids is shown at the top of the figure. Although knockdown of Panx and p15 weakened the repression by the forced tethering of Nxf2, the effects of H1- and HP1a-KD on the λN-Nxf2-mediated silencing were negligible, which is consistent with the results from Figures 4D and S3F.

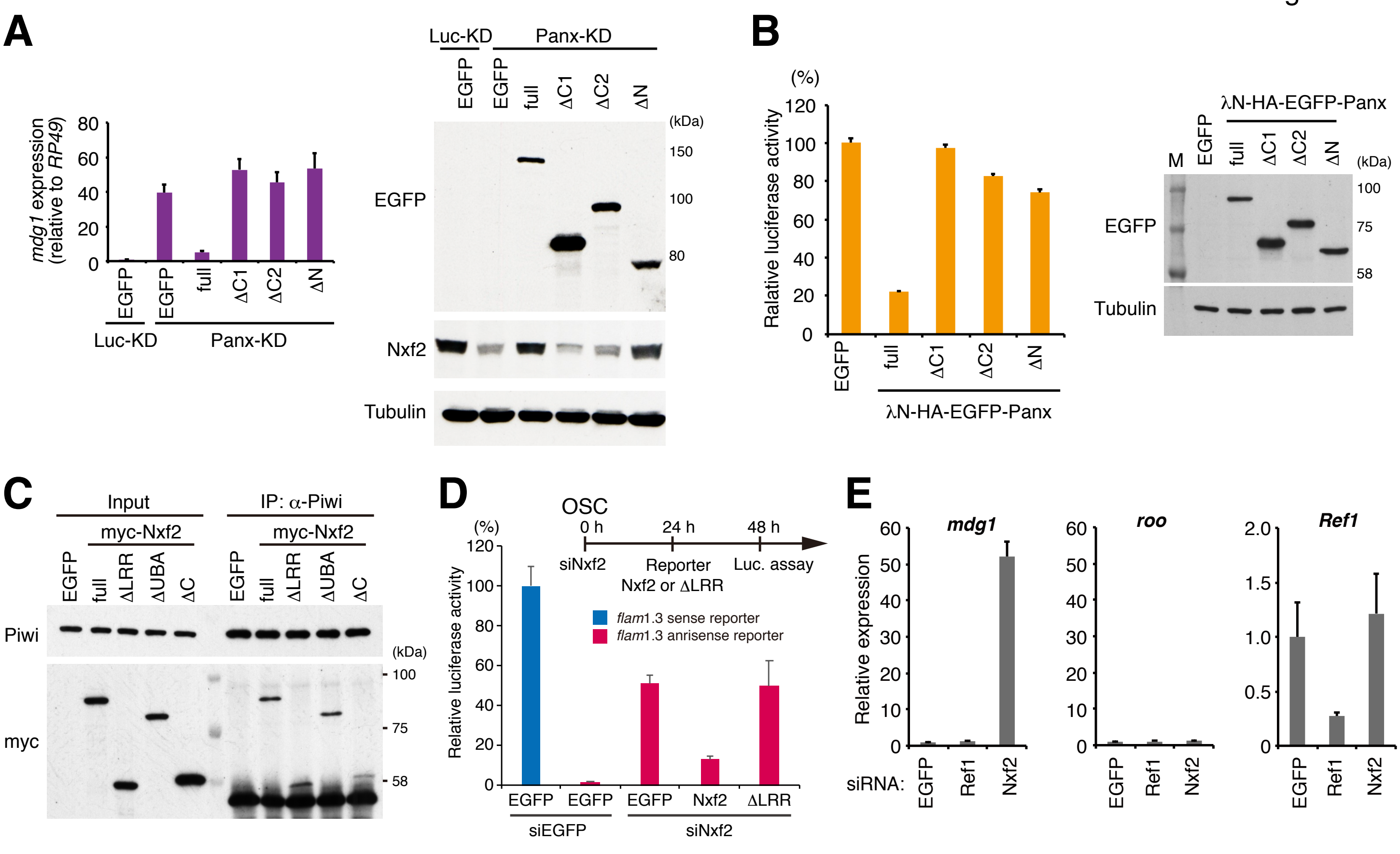

**Figure S5. Related to Figure 5**.

**(A)** EGFP-tagged Panx deletion constructs were expressed in Luc- (control) or Panx-depleted OSCs. *mdg1* expression levels were monitored by qRT-PCR. Expression values are normalised by the expression of *RP49*. Error bars indicate SD (n=3) (left panel). Western blotting (WB) with an antibody against EGFP, Nxf2, and Tubulin using lysates from transfected OSCs (right panel). The deletion mutants that cannot interact with Nxf2 or Panx could not induce silencing of TE. In addition, the C-terminal region of Panx (400–541 aa) is required for silencing of TE and stable expression of Nxf2.

**(B)** Luciferase fluorescence level in OSCs expressing boxB reporter luciferase and the indicated λN fusions (Panx deletion mutants). Error bars indicate SD (n=4) (left panel). WB of lysates from OSCs expressing boxB reporter luciferase and the indicated λN fusions using EGFP and tubulin antibody (right panel). The deletion mutants that cannot interact with Nxf2 or Panx could not induce silencing upon recruitment to reporter RNA.

**(C)** Immunoprecipitation (IP) from lysate of OSCs expressing myc-tagged Nxf2 proteins using anti-Piwi antibody, followed by WB using anti-myc and -Piwi antibodies. All mutant proteins of Nxf2 can interact with Piwi in OSCs.

**(D)** Silencing of reporter gene harboring *flam* element (1.3 kbp, antisense) occurs in an Nxf2-dependent manner. Transfection schedule of siRNA and plasmids is shown at the top of the figure. Exogenous expression of Nxf2 protein repressed reporter gene activity in OSCs, but that of Nxf2-ΔLRR protein did not. Error bar indicates SD (n=3).

**(E)** RNA levels of *mdg1*, *roo*, and *Ref1* were quantified by qRT-PCR upon depletion of EGFP (control), Ref1, or Nxf2. Expression levels are normalized by the expression of *RP49*. Error bars represent SD (n=3). Although Ref1 knockdown significantly decreased the expression level of *Ref1*, *mdg1* TE was unaffected.

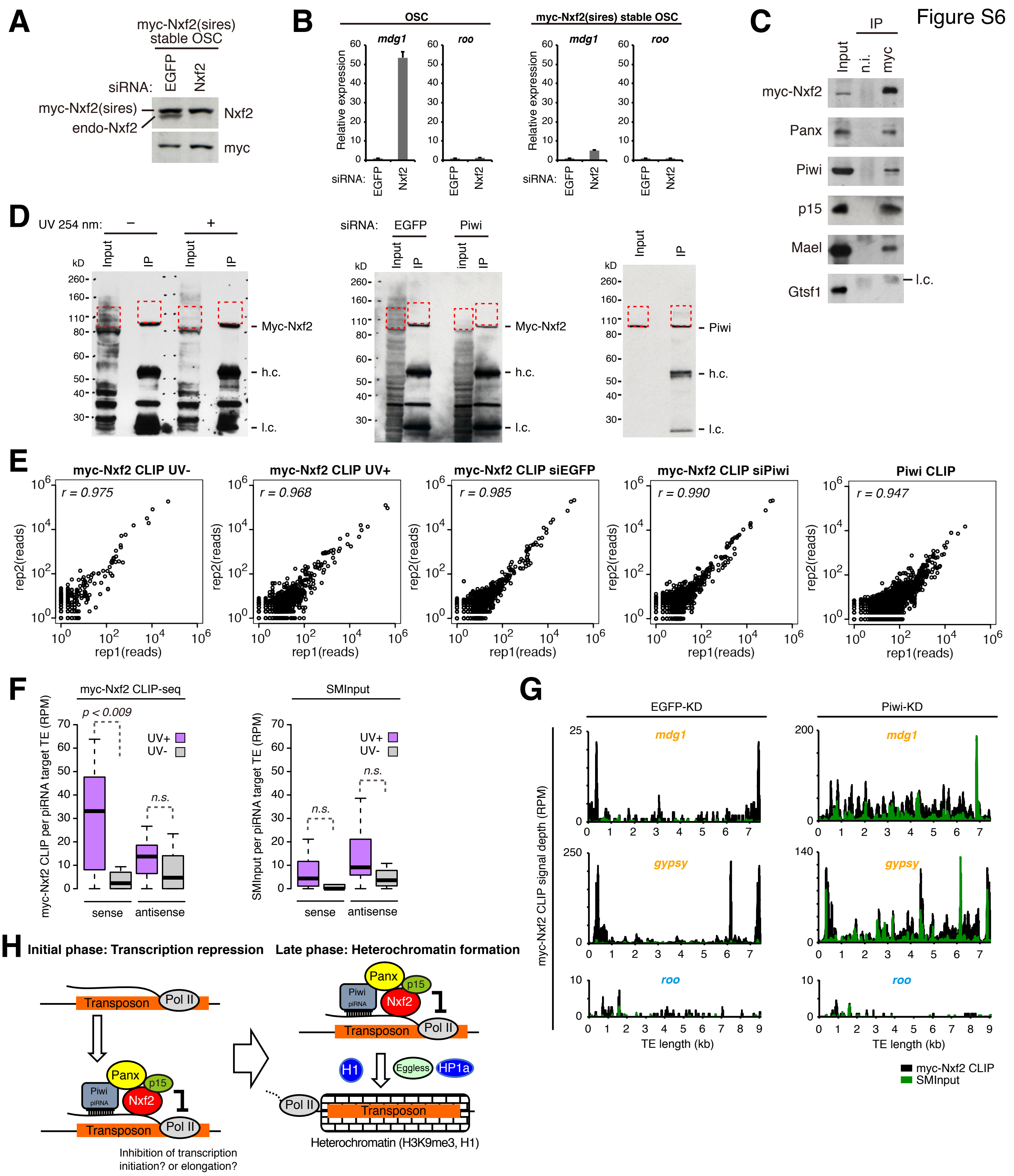

#### Figure S6. Related to Figure 6.

**(A)** Western blotting (WB) of myc-Nxf2 stable OSC lysate, using Nxf2 and myc antibody upon knockdown of EGFP (control) or Nxf2. Note that expressed myc-Nxf2 is resistant to siRNA.

**(B)** RNA levels of *mdg1* and *roo* were quantified by qRT-PCR upon depletion of EGFP (control) or Nxf2. Expression levels are normalized by the expression of *RP49*. Error bars represent SD (n=3). In OSCs without myc-Nxf2 expression, *mdg1* is de-silenced upon Nxf2-KD, whereas in the myc-Nxf2-expressing stable line, *mdg1* remains silenced.

**(C)** Immunoprecipitation (IP) from myc-Nxf2-expressing OSC lysate using anti-myc antibody, followed by WB of myc-Nxf2 (by myc antibody), Panx, Piwi, p15, Mael, and Gtsf1. l.c.: light chain from the antibody.

*(legend continued on next page)*

#### Figure S6. Related to Figure 6. (continued)

**(D)** WB of myc-Nxf2 IP during CLIP from non-irradiated or UV-irradiated cells (left panel). WB of myc-Nxf2 IP during CLIP from EGFP or Piwi knockdown cells (middle panel). WB of Piwi IP during CLIP (right panel). Red dotted line indicates the region excised for CLIP library preparation.

**(E)** Scatter plots of obtained CLIP reads in two biological replicates of indicated CLIP experiments. Each dot represents the read count of the peak called using the Piranha peak-calling algorithm (Uren et al., 2012).

**(F)** Boxplots showing piRNA-targeted TE-mapped read counts (RPM) obtained from myc-Nxf2 CLIP or SMInput, based on CLIP-seq performed on UV-irradiated and non-irradiated OSCs. Reads mapped in sense and antisense directions were calculated separately. Boxplot whiskers show maxima and minima. *P*-values were calculated by Wilcoxon rank-sum test.

**(G)** Density plots for myc-Nxf2 CLIP signal depth over the consensus sequence from *mdg1*, *gypsy* (targeted by Piwi–piRNA, in orange letters), and *roo* (not targeted by Piwi–piRNA, in blue letters) TEs in EGFP-, Piwi-KD OSCs. Reads obtained in myc-Nxf2 CLIP samples are indicated in black, where SMInput is indicated in green.

**(H)** Schematic model showing the two-phase regulation of TEs by Piwi–piRISC. Nxf2 forms a complex with Panx, Piwi, and p15, and associates with nascent RNA of target transposable elements. This complex regulates transcription of the TE by the inhibition of Pol II (initial phase). The co-transcriptionally regulated TE shifts to heterochromatin formation, mediated by H3K9me3 marks and H1 (late phase).

Figure S7

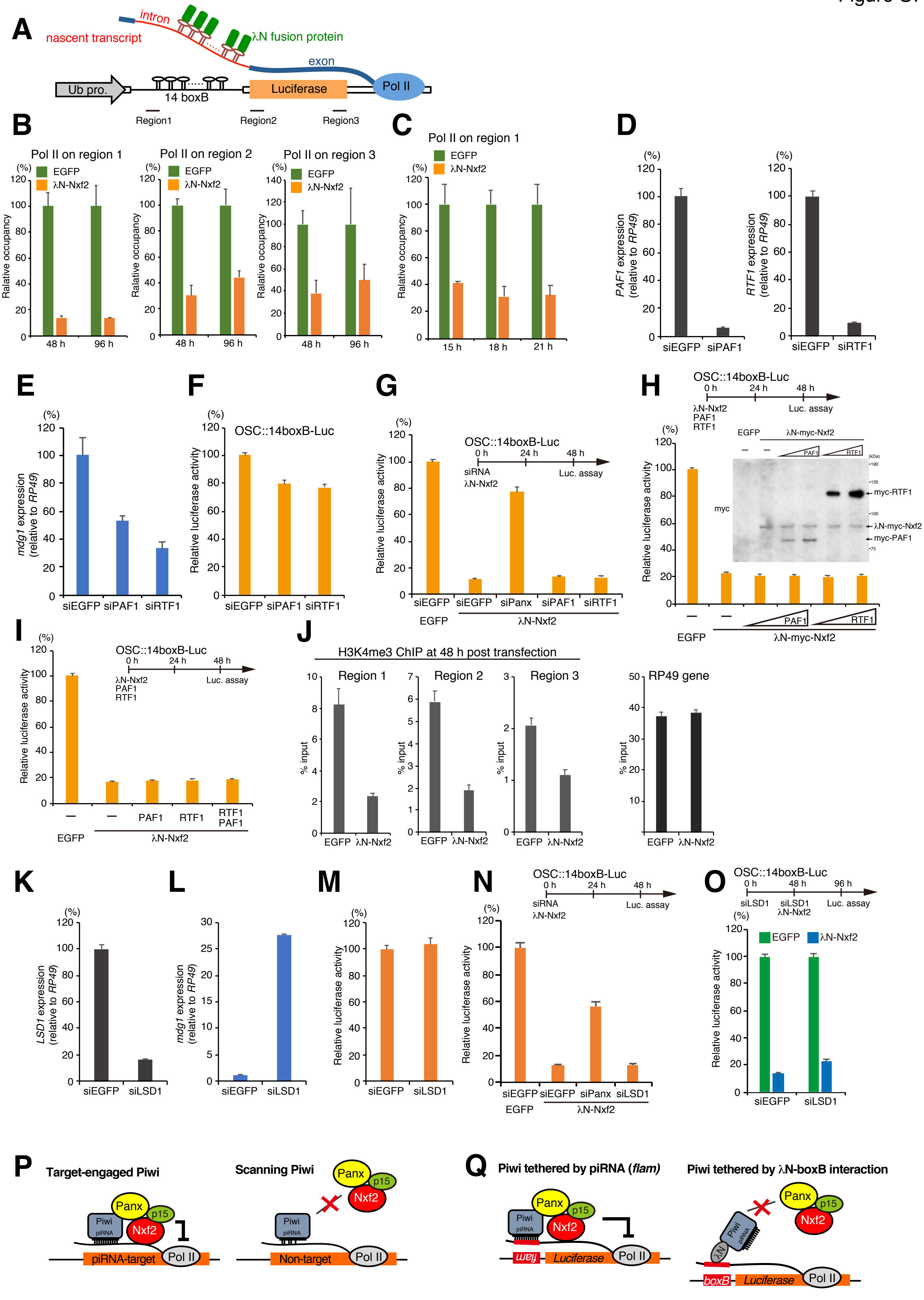

A
nascent transcript
intron
λN fusion protein
exon
Ub pro.
14 boxB
Luciferase
Pol II
Region1
Region2
Region3
B
Pol II on region 1
Pol II on region 2
Pol II on region 3
EGFP
λN-Nxf2
Ralative occupancy
48 h
96 h
C
Pol II on region 1
15 h
18 h
21 h
D
PAF1 expression (relative to RP49)
RTF1 expression (relative to RP49)
siEGFP
siPAF1
siRTF1
E
mdg1 expression (relative to RP49)
F
OSC::14boxB-Luc
Relative luciferase activity
G
OSC::14boxB-Luc
0 h
24 h
48 h
siRNA
λN-Nxf2
Luc. assay
siPanx
EGFP
H
OSC::14boxB-Luc
λN-Nxf2
PAF1
RTF1
λN-myc-Nxf2
myc
myc-RTF1
λN-myc-Nxf2
myc-PAF1
I
OSC::14boxB-Luc
RTF1 PAF1
J
H3K4me3 ChIP at 48 h post transfection
Region 1
Region 2
Region 3
RP49 gene
% input
K
LSD1 expression (relative to RP49)
siLSD1
L
mdg1 expression (relative to RP49)
M
N
OSC::14boxB-Luc
O
OSC::14boxB-Luc
0 h
48 h
96 h
siLSD1
siLSD1 λN-Nxf2
Luc. assay
P
Target-engaged Piwi
Scanning Piwi
Panx
p15
Nxf2
Piwi
piRNA
piRNA-target
Non-target
Pol II
Q
Piwi tethered by piRNA (flam)
Piwi tethered by λN-boxB interaction
flam
Luciferase
boxB
λN

#### Figure S7. Related to Discussion.

**(A)** The scheme shows regions targeted by qPCR, following ChIP experiments.

**(B)** ChIP-qPCR analysis of RNA polymerase II (Pol II) occupancy on the reporter gene upon λN-Nxf2 tethering. Bar graph shows the occupancy relative to that of EGFP. Error bars indicate SD (n=3). OSCs were harvested at 48 or 96 hpt.

**(C)** Time course of Pol II occupancy on region 1, immediate upstream of 14 boxB sites, upon λN-Nxf2 tethering. OSCs were harvested at the indicated time points. Error bars indicate SD (n=3). Contrary to our expectations, Pol II accumulation by the enforced tethering of Nxf2 at 14 boxB sites was not observed.

**(D)** RNA levels of *PAF1* and *RTF1* were quantified by qRT-PCR upon treatment of siEGFP (control), siPAF1, or siRTF1. Error bars represent SD (n=3).

**(E)** RNA level of *mdg1* was quantified by qRT-PCR upon treatment of siEGFP (control), siPAF1, or siRTF1. Error bars represent SD (n=3). Depletion of PAF1 or RTF1 enhanced the silencing of *mdg1* in OSCs.

**(F)** Depletion of PAF1 and RTF1 slightly weakened the luciferase activities from the reporter gene with 14 boxB sites integrated in the OSC genome. Error bars represent SD (n=4).

**(G)** Effects of knockdown of PAF1 and RTF1 on boxB reporter activity upon λN-Nxf2 expression. Bar graph shows luciferase activity relative to that of the sample co-transfected with myc-EGFP and siEGFP (control). Error bars indicate SD (n=4). Transfection schedule of siRNA and protein expression plasmids is shown at the top of the figures. Depletion of PAF1 or RTF1 had no impact on the silencing by the enforced tethering of Nxf2 on a nascent transcript.

**(H and I)** Effects of exogenous expression of PAF1 and RTF1 on boxB reporter activity upon λN-Nxf2 expression. Bar graph shows luciferase activity relative to that of the sample transfected with myc-EGFP (control). Error bars indicate SD (n=4). Transfection schedule of plasmids is shown at the top of the figures. Exogenous expression of PAF1 or RTF1 did not cancel the silencing by the enforced tethering of Nxf2 on a nascent transcript.

**(J)** ChIP-qPCR analysis of H3K4me3 occupancy on the reporter gene (14 boxB-Luc reporter gene integrated into the genome) upon λN-Nxf2 tethering. Bar graph shows % input. Error bars indicate SD (n=3). Regions targeted by qPCR are shown in (A). OSCs were harvested at 48 h post-transfection (hpt). H3K4 methylation is a transcriptionally active marker. Although occupancy of H3K4me3 on the *RP49* gene was not affected by λN-Nxf2 expression, the occupancy of H3K4me3 on the reporter gene decreased even at an earlier time point (48 hpt).

**(K)** RNA level of *LSD1* was quantified by qRT-PCR upon treatment of siEGFP (control) or siLSD1. Error bars represent SD (n=3).

**(L)** RNA level of *mdg1* was quantified by qRT-PCR upon treatment of siEGFP (control) or siLSD1. Error bars represent SD (n=3). mdg1 was de-silenced by LSD1 depletion.

**(M)** Luciferase activity was not affected by depletion of LSD1. Error bars represent SD (n=4)

**(N and O)** Effects of knockdown of LSD1 on boxB reporter activity upon λN-Nxf2 expression. Bar graph shows luciferase activity relative to that of the sample co-transfected with myc-EGFP and siEGFP (control). Error bars indicate SD (n=4). Transfection schedule of siRNA and protein expression plasmids is shown at the top of the figures. Depletion of LSD1 had limited impact on the silencing by the forced tethering of Nxf2 on a nascent transcript.

**(P)** Schematic model showing the target-engaged Piwi. "The target-engaged Piwi" can recruit the Panx-Nxf2-p15 complex to the target, which elicits co-transcriptional repression. However, "the scanning Piwi" is in the process of searching for targets through random interaction between piRNA and transcripts, many of which likely show partial complementarity. The scanning Piwi cannot recruit Panx-Nxf2-p15 complex to the target.

**(Q)** Schematic model showing Piwi that is targeted by piRNA or is tethered by λN-boxB interaction in the reporter systems. Piwi targeted by piRNA derived from the *flam* locus can recruit the Panx-Nxf2-p15 complex to the reporter gene, leading to co-transcriptional silencing. Piwi tethered by λN-boxB interaction cannot recruit the Panx-Nxf2-p15 complex to the reporter gene, and thus fails to induce silencing.

Upper band

| Rank | Description | Accession | emPAI | Score | Mass | Sequence coverage (%) |
|---|---|---|---|---|---|---|
| 1 | Piwi [Drosophila melanogaster] | 429892668 | 8.06 | 1054 | 97037 | 70 |
| 2 | piwi, partial [Drosophila simulans] | 418207352 | 2.65 | 535 | 95840 | 46 |
| 3 | GH10652p [Drosophila melanogaster] | 20976828 | 2.05 | 698 | 88155 | 51 |
| 4 | rasputin [Drosophila melanogaster] | 7739653 | 1.63 | 531 | 74940 | 43 |
| 5 | DNA ligase III [Drosophila melanogaster] | 84796152 | 1.33 | 579 | 90791 | 44 |
| 6 | Fmr1, isoform G [Drosophila melanogaster] | 383292605 | 1.32 | 303 | 80980 | 29 |
| 7 | CG10077, partial [Drosophila busckii] | 924558740 | 0.72 | 294 | 86515 | 30 |
| 8 | combgap, isoform B [Drosophila melanogaster] | 21645407 | 0.67 | 270 | 83703 | 14 |
| 9 | single-stranded recognition protein [Drosophila melanogaster] | 290280 | 0.6 | 227 | 81645 | 30 |
| 10 | belle, isoform A [Drosophila melanogaster] | 17985987 | 0.57 | 162 | 85029 | 24 |

Middle band

| Rank | Description | Accession | emPAI | Score | Mass | Sequence coverage (%) |
|---|---|---|---|---|---|---|
| 1 | Fmr1, isoform A [Drosophila melanogaster] | 19922726 | 7.94 | 1237 | 76030 | 64 |
| 2 | hsp 82 [Drosophila melanogaster] | 8127 | 4.9 | 946 | 81814 | 61 |
| 3 | nuclear RNA export factor 2 (NXF2) [Drosophila melanogaster] | 14456090 | 2.03 | 703 | 96573 | 41 |
| 4 | Nnp-1 [Drosophila melanogaster] | 7298046 | 1.02 | 276 | 78947 | 34 |
| 5 | belle, isoform A [Drosophila melanogaster] | 7299061 | 0.83 | 201 | 85029 | 32 |
| 6 | Bj6 protein [Drosophila melanogaster] | 7662 | 0.74 | 201 | 77005 | 27 |
| 7 | DNA replication factor MCM7 [Drosophila melanogaster] | 4903288 | 0.69 | 155 | 81232 | 22 |
| 8 | eIF3-S9, isoform A [Drosophila melanogaster] | 7302767 | 0.61 | 157 | 80391 | 30 |
| 9 | GH14068 [Drosophila grimshawi] | 193891778 | 0.53 | 322 | 80175 | 19 |
| 10 | rasputin [Drosophila melanogaster] | 7739653 | 0.41 | 110 | 74940 | 14 |

Lower band

| Rank | Description | Accession | emPAI | Score | Mass | Sequence coverage (%) |
|---|---|---|---|---|---|---|
| 1 | CG9754/Panoramix/Silencio [Drosophila melanogaster] | 7291281 | 5.57 | 617 | 61110 | 45 |
| 2 | no-on-transient A product form I [Drosophila melanogaster] | 157978 | 4 | 610 | 76920 | 48 |
| 3 | AT19081p [Drosophila melanogaster] | 21429758 | 2.82 | 499 | 79722 | 44 |
| 4 | RecName: Full=Protein claret segregational | 127945 | 2.75 | 684 | 77426 | 55 |
| 5 | glucosidase 2 beta subunit, isoform A [Drosophila melanogaster] | 7298396 | 1.46 | 309 | 61501 | 34 |
| 6 | uncharacterized protein Dmel_CG10565, isoform A [Drosophila melanogaster] | 7296390 | 0.79 | 150 | 73575 | 23 |
| 7 | Fragile X related [Drosophila melanogaster] | 6503206 | 0.76 | 208 | 75773 | 28 |
| 8 | belle, isoform A [Drosophila melanogaster] | 7299061 | 0.65 | 134 | 85029 | 32 |
| 9 | CG8368, isoform A [Drosophila melanogaster] | 7295335 | 0.56 | 183 | 76671 | 35 |
| 10 | LD25239p [Drosophila melanogaster] | 16198023 | 0.52 | 196 | 71051 | 10 |

**Table S1. List of proteins detected in MS analysis (related to Figure 1)**

Proteins detected by MS analysis of indicated bands from Panx-immunoprecipitation followed by silver staining (Figure 1A) are listed with their accession number, emPAI, Score, Mass, and Sequence coverage.

| Protein | Annotation Symbol | Significance (p-value, -log10) | Enrichment (peptide count in flag-Nxf2 IP +1/Control IP +1, log2) |
|---|---|---|---|
| Nxf2 | CG4118 | 5.32 | 5.71 |
| Hsc70-4 | CG4264 | 4.05 | 5.23 |
| betaTub56D | CG9277 | 3.50 | 4.52 |
| Panx | CG9754 | 3.37 | 5.09 |
| alphaTub84B\|alphaTub84D\|alphaTub85E | CG1913 | 3.36 | 4.86 |
| HtrA2 | CG8464 | 3.35 | 4.17 |
| betaTub56D\|betaTub85D | CG9277 | 2.92 | 3.91 |
| alphaTub84B\|alphaTub84D | CG1913 | 2.52 | 4.17 |
| blw | CG3612 | 2.49 | 3.46 |
| Hsc70-3\|Hsc70-4 | CG4147 | 2.49 | 3.46 |
| CG7194 | CG7194 | 2.22 | 4.46 |
| CkIIalpha | CG17520 | 2.22 | 3.00 |
| Hrb98DE | CG9983 | 2.22 | 3.00 |
| Ant2\|sesB | CG1683 | 1.98 | 3.58 |
| Droj2 | CG8863 | 1.91 | 3.91 |
| DnaJ-1 | CG10578 | 1.86 | 2.81 |
| betaTub56D\|betaTub60D\|betaTub85D | CG9277 | 1.81 | 2.58 |
| Cctgamma | CG8977 | 1.81 | 2.58 |
| p15 (Nxt1) | CG12752 | 1.79 | 3.17 |
| UQCR-C2 | CG4169 | 1.74 | 4.25 |
| Fib | CG9888 | 1.74 | 3.32 |
| Ssrp | CG4817 | 1.72 | 3.58 |
| ATPsynbeta | CG11154 | 1.62 | 3.70 |
| lost | CG14648 | 1.55 | 4.46 |
| porin | CG6647 | 1.52 | 3.46 |
| Ef1alpha100E\|Ef1alpha48D | CG1873 | 1.49 | 4.25 |
| Hsp68 | CG5436 | 1.45 | 3.00 |
| sqd | CG16901 | 1.45 | 4.00 |
| T-cp1 | CG5374 | 1.45 | 4.00 |
| Hsp26 | CG4183 | 1.39 | 3.17 |
| Rpn2 | CG11888 | 1.24 | 3.32 |
| Grip71 | CG10346 | 1.20 | 3.46 |
| CG14207 | CG14207 | 1.10 | 2.58 |
| Rpt6 | CG1489 | 1.10 | 2.58 |
| AGO2 | CG7439 | 1.08 | 4.17 |
| Hsp83 | CG1242 | 1.06 | 4.32 |
| Hsc70-3 | CG4147 | 0.99 | 2.81 |
| l(2)gl | CG2671 | 0.90 | 3.91 |
| CG6439 | CG6439 | 0.88 | 3.00 |
| Hsp23 | CG4463 | 0.87 | 3.32 |
| Hsp27 | CG4466 | 0.85 | 2.58 |
| hyd | CG9484 | 0.82 | 4.39 |
| Tcp-1zeta | CG8231 | 0.80 | 4.25 |
| CG9281 | CG9281 | 0.74 | 4.39 |
| dre4 | CG1828 | 0.74 | 3.17 |
| eIF-4a | CG9075 | 0.74 | 3.17 |
| spn-E | CG3158 | 0.74 | 2.81 |
| pAbp | CG5119 | 0.73 | 4.09 |
| Piwi | CG6122 | 0.71 | 2.58 |
| 14-3-3zeta | CG17870 | 0.71 | 2.58 |
| bel | CG9748 | 0.71 | 2.58 |
| Hop | CG2720 | 0.71 | 2.58 |
| Nap1 | CG5330 | 0.71 | 2.58 |
| PyK | CG7070 | 0.71 | 2.58 |
| r | CG18572 | 0.71 | 2.58 |
| Rpt3 | CG16916 | 0.71 | 2.58 |
| Mtor | CG8274 | 0.70 | 3.81 |
| Dcr-2 | CG6493 | 0.70 | 3.17 |
| CG8258 | CG8258 | 0.56 | 2.58 |
| smid | CG8571 | 0.56 | 2.58 |

**Table S2. List of proteins detected in shotgun proteome analysis of flag-Nxf2 (related to Figure 2)**

Proteins detected with shotgun proteome analysis of flag-Nxf2 immunoprecipitants are listed. Enrichment of peptide obtained by flag-Nxf2 IP over Control IP (log2 value of peptide count+1), and significance calculated by replicate experiments as the p-value (-log10) are listed along with protein name and annotation symbol. Proteins with significance over 0.5 and enrichment over 2.5 are indicated (ribosomal proteins are excluded from the list).

### Table S3

| Name | Sequence | Usage | Vector construction method |
|---|---|---|---|
| BoxB_EcoRIfor | gatGAATTCGCTAGCTTCCCTAAGTCCAACT | Described in the Methods section | Described in the Methods section |
| BoxB_XhoIrev | gacCTCGAGATAATATCCTCGATAGGGCC | | |
| BoxB_SalIfor | gatGTCGACGCTAGCTTCCCTAAGTCCAACT | | |
| Fluc_KpnIfor | acgtggtaccaacatggccgaagacgccaaaaacataaagaaag | pAc-Fluc-5boxB vector construction | RE digest with KpnI/EcoRI |
| Fluc_EcoRIrev | acgtgaattcttaCAATTTGGACTTTCCGCCCTTCTTGG | | |
| Ubpro_BamHIfor | GATggatccGAAGCGACTGGCGATTCGCGTG | Described in the Methods section | Described in the Methods section |
| Ubpro_KpnIrev | ACGTggtaccGGATTATTCTGCGGGAAGAAAATAGAGATGTG | | |
| lambdaN_KpnIfor | GTTCATTGGTACCCAACATGGACGCA | pUASp-lambdaN-MCS vector construction | RE digest with KpnI/NotI |
| lambdaN_NotIrev | ctcgagcggccgcgaattcggatccGCTAGCgcGTAATCTGGAACATC GTATGGGTAAGCC | | |
| lamdaN_NheI_for | acgtGCTAGCaacATGGACGCACAAACACGACGACGTG | Described in the Methods section | Described in the Methods section |
| lamdaN_NheI_rev | acgtGCTAGCgcAGCGTAATCTGGAACATCGTATGGGTAAG | | |
| Nxf2_HindIIIfor | ATAAGCTTATGCCAAACCAGATGAGAG | pAcM-Nxf2 vector construction | RE digest with HindIII/EcoRI |
| Nxf2_EcoRIrev | ATGAATTCCTAGGCGAATGCTAGATC | | |
| Nxf2-sires-INVfor | GAACGAGATAACTGTCCATCATAAGGGCCGCCTGGAA | pAcM-Nxf2_sires vector construction | inverse PCR from pAcM-Nxf2 |
| Nxf2-sires-INVrev | CTTATGATGGACAGTTATCTCGTTCCAATGGGTACCC | | |
| Nxf2delLRR_HindIIIfor | ATAAGCTTAACATGCTGCACAAGATCTTT | pAcM-Nxf2-ΔLRR vector construction | RE digest with HindIII/EcoRI |
| Nxf2delLRR_EcoRIrev | ATGAATTCCTAGGCGAATGCTAGATC | | |
| Nxf2delUBA_HindIIIfor | ATAAGCTTATGCCAAACCAGATGAGAG | pAcM-Nxf2-ΔUBA vector construction | RE digest with HindIII/EcoRI |
| Nxf2delUBA_EcoRIrev | ATGAATTCTTAATCCGGAAAATTTACACTGAAAG | | |
| Nxf2delC-HindIIIfor | ATAAGCTTATGCCAAACCAGATGAGAG | pAcM-Nxf2-ΔC vector construction | RE digest with HindIII/EcoRI |
| Nxf2delC_EcoRIrev | ATGAATTCTTACAACGACTGGCCAGGATT | | |
| Nxf2delNTF2_INVfor | CCAAGTGAACGTCAGGCGGA | pAcM-Nxf2-ΔNTF2 vector construction | inverse PCR from pAcM-Nxf2-sires |
| Nxf2delNTF2_INVrev | CAACGACTGGCCAGGATTCG | | |
| Nxf1_HindIIIfor | ATAAGCTTATGCCCAAACGCGGCGGT | pAcM-Nxf1 vector construction | RE digest with HindIII/EcoRI |
| Nxf1_EcoRIrev | ATGAATTCTTACTTCATAAAAGCCTCAGGCGG | | |
| Nxf3_HindIIIfor | ATAAGCTTATGGGATCCGTGCTGGAC | pAcM-Nxf3 vector construction | RE digest with HindIII/EcoRI |
| Nxf3_EcoRIrev | ATGAATTCTTAAAGGTCATCGCATCCAAACAGG | | |
| Panx_KpnIfor | AATGGTACCATGGAAGCTCCGATGAAGCT | pAcEGFP-Panx vector construction | RE digest with KpnI/NotI |
| Panx_NotIrev | ATTGCGGCCGCTCACTATGGCTGTCGACCCTTTA | | |
| Panx-sires_INVfor | CCTAAGATGGAACCGAAGATCAAAGAAGATGCCG | pAcEGFP-Panx_sires vector construction | inverse PCR from pAcEGFP-Panx |
| Panx-sires_INVrev | CGGCATCTTCTTTGATCTTCGGTTCCATCTTAGG | | |
| PanxdelC1_INVfor | TAGGCGGCCGCTCGAGTCTA | pAcEGFP-Panx-ΔC1 vector construction | inverse PCR from pAcEGFP-Panx-sires |
| PanxdelC1_INVrev | ATTGGAGAATTTAATAGGTC | | |
| PanxdelC2_INVfor | TAGGCGGCCGCTCGAGTCTA | pAcEGFP-Panx-ΔC2 vector construction | inverse PCR from pAcEGFP-Panx-sires |
| PanxdelC2_INVrev | CAAGCCCCATTTAGACAAAT | | |
| PanxdelN_INVfor | CCGCAAAATGATACGCCCAT | pAcEGFP-Panx-ΔN vector construction | inverse PCR from pAcEGFP-Panx-sires |
| PanxdelN_INVrev | CATGGTACCGTCGACTGCAG | | |
| p15_HindIIIfor | AATAAGCTTATGGACAGCGATTTGAAAGC | pAcM-p15 vector construction | RE digest with HindIII/NotI |
| p15_NotIrev | AATGCGGCCGCTCAGACCTCCTGCATTCGGT | | |
| p15-sires_INVfor | CAGCAGATAGGTCGCCTCTATTTGG | pAcM-p15_sires vector construction | inverse PCR from pAcM-p15 |
| p15-sires_INVrev | TCTCCTATTGTCCACGGAGGCGTAG | | |
| Panx-mid_BamHIfor | AATGGATCCCCGCAGCTGATCTCATTGCACA | pGEX-Panx-mid vector construction | RE digest with BamHI/NotI |
| Panx-mid_NotIrev | AATGCGGCCGCTCAGGACTCATTAACAGGGAGTT | | |
| Nxf2Full-EcoRIfor | atgaattcATGCCAAACCAGATGAGAG | pPET-Nxf2-full vector construction | RE digest with EcoRI/SalI |
| Nxf2Full-SalIrev | ATGTCGACCTAGGCGAATGCTAGATC | | |
| Nxf2LRR_BamHIfor | ATTGGATCCCCATGCCAAACCAGATGAGAG | pGEX-Nxf2-LRR vector construction | RE digest with BamHI/NotI |
| Nxf2LRR_NotIrev | ATGCGGCCGCCTAGAAAGGCTTTCCATCCA | | |
| p15_BamHIfor | ATTGGATCCCCATGGACAGCGATTTGAAAG | pGEX-p15 vector construction | RE digest with BamHI/NotI |
| p15_NotIrev | AATGCGGCCGCTCAGACCTCCTGCATTCGGT | | |
| egg_INVfor | TGCGAATTGTCGCTCCAAGT | pGEX-Egg-Nter vector construction | inverse PCR from pGEX-Egg-full |
| egg_INVrev | TAAGCGGCCGCATCGT | | |
| PAF1for_KpnI | acgtggtaccATGCCACCCACGATCAACAATTCGGC | pAcM-PAF1 vector construction | RE digest with KpnI/NotI |
| PAF1rev_NotI | ACGTgcggccgcTCATTCATCTGAGGCGCTTCCAGAGC | | |
| RTF1for_KpnI | acgtggtaccATGGGCAAACGGCGGACGCAATC | pAcM-RTF1 vector construction | RE digest with KpnI/NotI |
| RTF1rev_NotI | ACGTgcggccgcCTAGATAAGACCGCGCTTTTTTTTGTAATCCT | | |
| Panx_for | CAAAGAGGAGCCACCGGAAT | Panx qPCR | |
| Panx_rev | CTGCGATGGCTTAACTGTGC | | |
| Nxf2_for | CCTCGAACGGTTTCACTCCA | Nxf2 qPCR | |
| Nxf2_rev | AGATCCTGCACCCAGTTGTG | | |
| Piwi_for | CGACCAAGAACCGTAGCCTT | Piwi qPCR | |
| Piwi_rev | CTCCGTGACTGTGCTGAAGT | | |
| mdg1_for | AACAGAAACGCCAGCAACAGC | mdg1 qPCR | |
| mdg1_rev | CGTTCCCATGTCCGTTGTGAT | | |
| roo_for | CGTCTGCAATGTACTGGCTCT | roo qPCR | |
| roo_rev | CGGCACTCCACTAACTTCTCC | | |
| HetA_for | CGCGCGGAACCCATCTTCAGA | HetA qPCR | |
| HetA_rev | CGCCGCAGTCGTTTGGTGAGT | | |
| Ref1_for | GCGCTTGGAAGCACGATATG | Ref1 qPCR | |
| Ref1_rev | CGTAGTCCAGGTTACCGACG | | |
| RP49_for | CCGCTTCAAGGGACAGTATCTG | RP49 qPCR | |
| RP49_rev | ATCTCGCCGCAGTAAACGC | | |
| 14BoxBLuc_ChIPfor | CAGCCACGCATCTCGTTTTC | boxB qPCR | |
| 14BoxBLuc_ChIPrev | AGGGAAGCTAGCGTCGAGTGCGT | | |
| Fluc_4_ChIPfor | GCCGAAGACGCCAAAAACAT | Fluc(4-300) qPCR | |
| Fluc_300_ChIPrev | TCCGATAAATAACGCGCCCA | | |
| Fluc_1277RTfor | ACTGGGACGAAGACGAACAC | Fluc(1277-1512) qPCR | |
| Fluc_1512RTrev | GGCGACGTAATCCACGATCT | | |
| PAF1_for | GCTGGAGGAGGAGACTCTGA | PAF1 qPCR | |
| PAF1_rev | TCTTGGTGTCGCTGAAGGTC | | |
| RTF1_for | CGCTGGCGTCTCTGATTACT | RTF1 qPCR | |
| RTF1_rev | TCACGGCCGATTTACAACCA | | |
| LSD1_for | TCCAATCTCGCCTGCCATTT | LSD1 qPCR | |
| LSD1_rev | ACTCGCCTTACCAAACTGGG | | |
| Piwi-siRNA_for | GCUCCCAGGCGUGAAGGUGtt | Piwi RNAi-KD | |
| Piwi-siRNA_rev | CACCUUCACGCCUGGGAGCtt | | |
| Panx-siRNA_for | CCGAAGAUGGAGCCCAAGAtt | Panx RNAi-KD | |
| Panx-siRNA_rev | UCUUGGGCUCCAUCUUCGGtt | | |
| Nxf2-siRNA_for | CGAAAUCACAGUGCACCAUtt | Nxf2 RNAi-KD | |
| Nxf2-siRNA_rev | AUGGUGCACUGUGAUUUCGtt | | |
| p15-siRNA_for | CCGACGCCAACAAATTGGAtt | p15 RNAi-KD | |
| p15-siRNA_rev | UCCAAUUUGUUGGCGUCGGtt | | |
| HP1a-siRNA_for | GAUCUUGGGUGCCUCCGACtt | HP1a RNAi-KD | |
| HP1a-siRNA_rev | GUCGGAGGCACCCAAGAUCtt | | |
| H1-siRNA_for | GAAAGAUCCAGAUGCACCCtt | H1 RNAi-KD | |
| H1-siRNA_rev | GGGUGCAUCUGGAUCUUUCtt | | |
| Mael-siRNA_for | CGCCAAGAUGUCCCAUGAUtt | Mael RNAi-KD | |
| Mael-siRNA_rev | AUCAUGGGACAUCUUGGCGtt | | |
| Gtsf1-siRNA_for | GCTCCAGCAGCACATCTTAtt | Gtsf1 RNAi-KD | |
| Gtsf1-siRNA_rev | UAAGAUGUGCUGCUGGAGCtt | | |
| Egg-siRNA_for | GGTCACAAGCGTATTAGCTtt | Egg RNAi-KD | |
| Egg-siRNA_rev | AGCUAAUACGCUUGUGACCtt | | |
| Ref1-siRNA-for | CAGCGCUUGGAAGCACGAUtt | Ref1 RANi-KD | |
| Ref1-siRNA-rev | AUCGUGCUUCCAAGCGCUGtt | | |
| EGFP-siRNA_for | GGCAAGCUGACCCUGAAGUtt | EGFP RNAi-KD | |
| EGFP-siRNA_rev | ACUUCAGGGUCAGCUUGCCtt | | |
| Luc-siRNA_for | CGUACGCGGAAUACUUCGAtt | Luc RNAi-KD | |
| Luc-siRNA_rev | UCGAAGUAUUCCGCGUACGtt | | |
| PAF1-siRNA_for | GCCACCGCUUCGUGCAGUAtt | PAF1 RNAi-KD | |
| PAF1-siRNA_rev | UACUGCACGAAGCGGUGGCtt | | |
| RTF-siRNA_for | GGACGAAGAUGACAUGCAAtt | RTF1 RNAi-KD | |
| RTF-siRNA_rev | UUGCAUGUCAUCUUCGUCCtt | | |
| LSD1-siRNA_for | GUCAGGAUCGGCAUCCAUGtt | LSD1 RNAi-KD | |
| LSD1-siRNA_rev | CAUGGAUGCCGAUCCUGACtt | | |
| endo-siRNA-sl1 | GGAGCGAACTTGTTGGAGTCAA | esiRNA sl1 northern blot | |
| tj-piRNA | GGTAATGGGAATGCACTTCTCTTGAA | tj-piRNA northern blot | |
| Idefix-piRNA | AAACTACTGGCAATCGTTTGGGAA | idefix-piRNA northern blot | |

**Table S3. DNA oligonucleotides and siRNAs used in this study, related to experimental procedures**

**Supplemental Experimental Procedures:**

**Cell culture and transfection**

Ovarian somatic cells (OSCs) were cultured in Shield and Sang M3 Insect Medium (Sigma) supplemented with 10% fly extract, 10% foetal bovine serum, 0.6 mg/ml glutathione, and 10 μg/ml insulin as described previously (Niki et al., 2006; Saito et al., 2009). For the transfection of small interfering (si)RNA into OSCs, 40–300 pmol siRNA duplex was nucleofected into $3.0 \times 10^6$ cells using the Cell Line 96-well Nucleofector Kit SF (Lonza) and program DG150 of the 96-well Shuttle Device (Lonza). The siRNAs were transfected twice for 4-day KD, and once for 2-day KD. The siRNA sequences are listed in *Table S3*. Co-transfection of siRNA and plasmid DNA was performed using the Cell Line Nucleofector Kit V (Lonza) and program T-029 of the Nucleofector II Device (Lonza). For immunoprecipitation, expression vectors were transfected using Xfect Transfection Reagent (TaKaRa Clontech), in accordance with the manufacturer's instructions.

***Drosophila* strains**

Oregon R (OR) was employed as a wild-type strain. Lines harbouring shRNA for Nxf2 (y[1] sc[*] v[1]; P{y[+t7.7] v[+t1.8]=TRiP.HMS00945}attP2/TM3, Sb[1]) and MTD-Gal4 lines (P{w[+mC]=otu-GAL4::VP16.R}1, w[*]; P{w[+mC]=GAL4-nos.NGT}40; P{w[+mC]=GAL4::VP16-nos.UTR}CG6325[MVD1]) were obtained from Bloomington Drosophila Stock Center (Stock# 33985 and 31777, respectively). For the tethering assay in fly ovary, GFP reporter line (If/CyO; tub>EGFP_5xBoxB_SV40 [attP2]/TM3, Ser), λN line as a control (pUASp>lambdaN-HA [attP40]/CyO; tub>EGFP_5xBoxB_SV40 [attP2]/TM3, Ser), λN-Piwi line (pUASp>lambdaN-HA-Piwi [attP40]/CyO; tub>EGFP_5xBoxB_SV40 [attP2]/TM3, Ser), and λN-Panx line (pUASp>lambdaN-HA-CG9754 [attP40]/CyO; tub>EGFP_5xBoxB_SV40 [attP2]/TM3, Ser) were obtained from Vienna Drosophila Resource Center (VDRC) (Stock# 313408, 313390, 313392, and 313393, respectively). All stocks were maintained at 25°C. For calculation of the egg hatching rate, 20 females were mated with Oregon R males in yeasted apple agar plates. The females were left in the plate to continue laying eggs after mating. Laid eggs were counted after 24 h and hatched eggs were counted after another 24 h.

**Construction of expression plasmids**

To produce expression vectors, full-length cDNA was amplified using primers described in *Table S3*. First-strand cDNA was synthesised from total RNA isolated from the *OR* ovaries or OSCs. PCR products were then inserted into the pGEX-5X expression vector (GE Healthcare Bioscience), pET28 expression vector (Novagen), pAcM vector(Saito et al., 2006), pAcF vector, or pAcEGFP vector (EGFP cloned into the myc tag region of the pAcM vector), using restriction enzyme cloning or In-Fusion HD Cloning Kit (Clontech) (described in *Table S3*). The pGEX-5X or pET28 expression vector was used to express GST- or HIS-fused proteins, and the pAcM, pAcF, or pAcEGFP vector was used to express myc-, Flag-, or EGFP-tagged proteins in OSCs. To yield siRNA-resistant mutants and mutants for domain mapping experiments, inverse PCR was performed using the described oligonucleotides (*Table S3*). The inserted fragment for the pAcF-Nxf2 vector was obtained by *Hind*III/*Eco*RI restriction enzyme digestion of the pAcM-Nxf2 vector and ligated into the pAcF vector. Note that plasmid construction for the tethering assay is described in the "Tethering assay using OSCs" and "Tethering assay in fly ovary" sections.

**Production of antibodies (anti-Panx, anti-Nxf2, anti-Eggless, anti-p15)**

The middle fragment of the Panx protein (181–361 aa), N-terminal region of the Eggless protein (1–405 aa), or full-length p15 protein (1–134 aa) fused with glutathione S-transferase (GST), and full-length

Nxf2 (1–841 aa) fused with histidine (HIS) were used as antigens to immunise mice. The anti-Panx, -Nxf2, and -Eggless monoclonal antibodies were produced essentially as described previously(Ishizuka et al., 2002; Saito et al., 2006). For the anti-p15 antibody, serum from immunised mice was used for a polyclonal antibody.

**Immunoprecipitation**

To obtain whole-cell lysate, OSCs were washed in PBS and lysed in IP buffer (50 mM HEPES-KOH pH 7.4, 200 mM KCl, 1 mM EDTA, 1% Triton X-100, 0.1% Na-deoxycholate or 30 mM HEPES-KOH pH 7.4, 150 mM KOAc, 5 mM MgOAc, 5 mM DTT, 0.1% NP40) followed by probe-sonication (BRANSON, SFX150, or Bioruptor). After centrifugation, the supernatant was used for immunoprecipitation. To prepare nuclear lysate, OSCs were washed in PBS and suspension buffer (10 mM HEPES-KOH pH 7.4, 10 mM KCl, 1.5 mM $MgCl_2$, 0.5 mM DTT), and then homogenised in suspension buffer by passing through a syringe with a 25G needle. After centrifugation, supernatant was recovered as cytoplasmic lysate. The nuclear pellet was washed once and then re-suspended in IP buffer, followed by probe-sonication. The nuclear lysate was centrifuged and the supernatant was kept for immunoprecipitation. Antibodies were immunised on Dynabeads Protein G (Thermo Fisher) and incubated with lysates for 1–2 h, at 4°C. Then, beads were washed three to five times in IP buffer. Control immunoprecipitation using mouse nonimmune IgG (IBL) was conducted in parallel.

**Western blotting**

Western blotting was performed as described previously(Saito et al., 2006). Besides the antibodies generated in this study, anti-Piwi(Saito et al., 2006) (hybridoma supernatant), anti-DmGTSF1/Arx(Ohtani et al., 2013) (hybridoma supernatant), anti-Mael(Sato et al., 2011) (hybridoma supernatant), anti-H1(Iwasaki et al., 2016) (hybridoma supernatant), anti-Tubulin (E7, hybridoma supernatant), anti-myc (9E10, hybridoma supernatant), anti-flag (M2, sigma) (1:1,000), anti-H3 (ab1791, Abcam) (1:2,000), anti-HP1a (C1A9, DSHB) (1:1,000), anti-HA (HA-7, Sigma) (1:1,000), and anti-GFP (11814460001, Roche) (1:1,000) antibodies were used.

**Silver staining and mass spectrometry analysis**

Silver staining was performed using the SilverQuest Silver Staining Kit (Invitrogen), in accordance with the manufacturer's instructions. The bands shown in Figure. 1A were excised, de-stained, and sent for liquid chromatography–tandem mass spectrometry (MS) analysis to identify the peptides (Support Center for Advanced Medical Sciences, The University of Tokushima, Japan). The results were analysed using Mascot (Matrix Science), and Scaffold4 (Proteome Software Inc.) was used to validate MS/MS-based peptides.

**Immunofluorescence**

Immunofluorescence of OSCs and *Drosophila* ovaries was performed using anti-Panx IgG1 (hybridoma supernatant) or anti-myc (9E10, hybridoma supernatant) antibody as described previously(Ishizuka et al., 2002). Alexa Fluor546- or 488-conjugated anti-mouse IgG1 (Molecular Probes) (1:1,000 dilution) was used as secondary antibody. Slides were mounted using VECTASHIELD Mounting Medium with DAPI (Vector Laboratories).

**Quantitative reverse-transcription PCR and northern blotting**

qRT-PCR analysis and northern blotting were performed as previously described(Iwasaki et al., 2016). Oligonucleotides used for qRT-PCR primers and northern blotting probes are described in *Table S3.*

Previously described primers(Sienski et al., 2015) were used for qRT-PCR analysis of TE expression levels in ovary.

**Shotgun proteome analysis**

Immunoprecipitated proteins were eluted with 100 mM Tris-HCl (pH 8) and 2% sodium dodecyl sulphate (SDS). To remove the SDS from the eluted samples, the methanol–chloroform protein precipitation method was used. Briefly, four volumes of methanol, one volume of chloroform, and three volumes of water were added to the eluted sample and mixed thoroughly. Samples were centrifuged at 20,000 g for 10 min, the water phase was carefully removed, and then four volumes of methanol were added to the samples, followed by centrifugation at 15,000 rpm for 10 min. The supernatant was removed, and the pellet was washed once with 100% ice-cold acetone. Precipitated proteins were re-dissolved in guanidine hydrochloride and reduced with Tris(2-carboxyethyl)phosphine hydrochloride, alkylated with iodoacetamide, and then digested with lysyl endopeptidase and trypsin. Digested peptides were analysed using a direct nanoflow liquid chromatography system coupled to a time-of-flight mass spectrometer (QSTAR Elite, Sciex). Mass spectrometry and tandem mass spectrometry spectra were obtained in the information-dependent acquisition mode and were queried against the NCBI non-redundant database with an in-house Mascot server(Natsume et al., 2002) (version 2.2.1; Matrix Science).

**EMSA**

For EMSA, 16-mer single-stranded RNA (5′-AGCACCGUAAAGACGC-3′) described previously(Vourekas et al., 2015) was 5′-end-radiolabelled by p32 and 1 nM was incubated with the indicated amount of GST protein for 15 min at room temperature in 10 µl of RNA-binding buffer [50 mM Tris-HCl (pH 7.5), 50 mM KOAc, 2 mM $MgCl_2$, 1 mM DTT, and 1 U/µl RNasin]. After incubation, 1 µl of 50 mg/ml heparin was added and incubated for another 10 min, and the reactions were analysed using 5% native PAGE (Tris-glycine gel) at 4°C. The results were visualised by phospho-imaging.

**Tethering assay using OSCs**

We established a tethering assay system in OSCs based on the system in fly ovary, which was reported by Hannon's group(Yu et al., 2015). To obtain firefly luciferase-reporter constructs, a 5× boxB fragment was amplified from fly genomic DNA (VDRC #313408) and cloned into an *Eco*RI and *Xho*I sites of pAc5.1B, followed by insertion of luciferase fragment amplified from pCMV-luc at a *Kpn*I and *Eco*RI site, resulting in pAc-Fluc-5boxB. To obtain pAc-Fluc-10boxB, PCR-amplified 5× boxB carrying a *Sal*I and *Xho*I site was inserted into a *Xho*I site of pAc-Fluc-5boxB. To produce pUb-Fluc-10boxB, we replaced the *actin 5c* promoter of pAc-Fluc-10boxB with a ubiquitin promoter at *Bgl*II and *Kpn*I sites. The ubiquitin promoter was amplified from fly genomic DNA. For the intronic boxB reporter construct, we first removed the *Xho*I site in pAc-Fluc and replaced its *actin 5c* promoter with the ubiquitin promoter. Subsequently, 5× boxB fragments were inserted twice into the *Xho*I site within the intron of the ubiquitin promoter. In this process, a reporter construct with 14× boxB in the intron, pUb-14boxB-Fluc, was incidentally obtained, and its sequence was confirmed. For λN-HA fusion protein expression constructs, a λN-HA fragment was amplified from a plasmid, pUASp-λN-Empty (kindly provided by Dr. Brennecke), and inserted into an *Nhe*I site of pAcM-based and pAc-EGFP-based expression plasmids. The oligonucleotides used for plasmid construction are described in *Table S3.*

Isolation of the OSC line carrying reporter gene was performed essentially as described previously(Sumiyoshi et al., 2016). OSCs were co-transfected with a reporter construct and a plasmid carrying a blasticidin-resistance gene, using ScreenFect A (Wako), followed by selection with 25 µg/ml blasticidin for 24 h. Subsequently, 1 × $10^5$ cells were passaged in a 6-cm dish and allowed to form colonies in medium supplemented with or without a lower concentration of blasticidin (4–8 μg/ml). Some colonies were isolated and selected using a luciferase assay.

For the tethering assay, 3 × $10^6$ OSCs were transfected with 5 μg of plasmid DNA using the Cell Line Nucleofector Kit V (Lonza) and Program T-029 of the Nucleofector II Device (Lonza) and passaged in four wells of a 24-well tissue culture plate. Cells were lysed in 150 μl of 1× Glo Lysis Buffer (Promega). Aliquots of cell lysates were used to measure firefly luciferase activity using ONE-Glo Ex Luciferase Assay system (Promega) and Cytation 5 (BioTek). Total protein concentration of cell lysates was measured by a Bradford assay (Bio-Rad).

**Tethering assay in fly ovary**

Based on previous reports, we carried out a tethering assay in fly ovary(Sienski et al., 2015; Yu et al., 2015). For the RNA-tethering system in fly ovary, a DNA fragment of myc-Nxf2 was inserted at *Nhe*I and *Eco*RI sites in pUASp-λN-MCS, resulting in pUASp-λN-HA-myc-Nxf2. To obtain pUASp-λN-MCS, a λN fragment including multiple cloning sites was inserted between *Kpn*I and *Not*I sites in pUASp-λN-Empty (kindly provided by Dr. Brennecke). Oligonucleotides used for plasmid construction are described in *Table S3*. A transgenic line was established by phiC31-mediated integration of pUASp-λN-HA-myc-Nxf2 into the attP40 landing site(Bischof et al., 2007; Markstein et al., 2008), resulting in a transgenic line (y'w1118;attp40{lambdaN-HA-myc-Nxf2}/CyO). Transgenic and GFP reporter flies were crossed with a double balancer (Sp/CyO; Pr/TM3, Ser, sb), and then the progeny were crossed together, resulting in a transgenic line carrying GFP reporter gene and λN-HA-myc-Nxf2 gene (y'w1118; attp40{lambdaN-HA-myc-Nxf2}/CyO; tub>EGFP_5xBoxB_SV40 [attP2]/TM3). Fly ovaries were fixed in 4% formaldehyde and stained with DAPI, followed by microscopic observation to detect GFP signal in the egg chamber.

**piRNA reporter gene assay**

We established a piRNA reporter assay system in OSCs, based on the reports by Lau's group (Post et al., 2014). We selected 0.6 kbp and 1.3 kbp *flam* elements, from which a number of piRNAs are derived, were cloned into the intron of pUb-Fluc in sense orientation or in antisense orientation relative to the *luciferase* transcripts. These *flam* elements were amplified using primers described in *Table S3*. DNA and siRNA transfection, and luciferase assays in OSCs and S2 cells were performed as described in the "Tethering assay using OSCs" section.

**ChIP-qPCR analysis**

For ChIP using anti-RNA polymerase II antibody (8WG16; Santa Cruz), OSCs were fixed in 0.3% formaldehyde (methanol-free; Thermo Fisher) and lysed in sonication buffer (50 mM Tris-HCl pH 7.9, 150 mM NaCl, 1 mM EDTA, 1% Triton X-100, 0.1% Na-deoxycholate), followed by probe-sonication (BRANSON, SFX150). After centrifugation, the supernatant was passed through a 0.22-μm syringe filter (Membrane Solutions) and used for immunoprecipitation. For ChIP using anti-H1(Iwasaki et al., 2016) (hybridoma supernatant), anti-H3K9me3 (0319, Active motif), and anti-H3K4me3 (ab8580, Abcam), OSCs were fixed in 1% formaldehyde and lysed in ChIP buffer (50 mM Tris-HCl pH 7.9, 5 mM EDTA, 0.1% SDS), followed by probe-sonication. After centrifugation, the supernatant was mixed with dilution buffer (50 mM Tris-HCl pH 7.9, 250 mM NaCl, 1.6% Triton X-100, 0.16% Na-deoxycholate), passed through a 0.22-μm syringe filter, and used for immunoprecipitation. Lysates were incubated with antibodies on Dynabeads-Protein G for 3 h, at 4°C. Beads were washed in a series of wash buffers:

low-salt wash buffer (20 mM Tris-HCl pH 7.9, 150 mM NaCl, 2 mM EDTA, 1% Triton X-100, 0.1% Na-deoxycholate), high-salt wash buffer (20 mM Tris-HCl pH 7.9, 500 mM NaCl, 2 mM EDTA, 1% Triton X-100, 0.1% Na-deoxycholate), LiCl wash buffer (10 mM Tris-HCl pH 7.9, 250 mM LiCl, 1 mM EDTA, 1% NP-40, 0.5% Na-deoxycholate), and TE (10 mM Tris-HCl pH 7.9, 1 mM EDTA). Protein–DNA complexes were eluted from beads in elute buffer (50 mM Tris-HCl pH 7.9, 10 mM EDTA, 1% SDS) at 65ºC for 10 min, followed by reversing cross-linking at 65ºC for over 12 h in the presence of 200 mM NaCl. DNA was recovered and quantified with the Dice Real Time System III (TaKaRa) using TB Green Fast qPCR Mix (TaKaRa) and primer sets for the reporter gene (described in *Table S3*).

**mRNA-seq analysis**

Preparation and sequencing of Poly(A)+ RNA libraries and computational analyses were performed as described previously(Iwasaki et al., 2016). Libraries were prepared from total RNA obtained from OSCs with 4-day knockdown of the indicated genes, and analysed by Illumina HiSeq. This yielded 10–17 million reads per sample. For computational analysis of the obtained reads, Bowtie 2.2.9(Langmead and Salzberg, 2012) was used with the default parameters to map sequences to the *Drosophila* genome (dm3). Reads mapped to the genome were then mapped to the *Drosophila melanogaster* TE consensus sequence from Repbase (Jurka, 1998), using Bowtie 1.0.1, and those that were uniquely mapped were extracted. Expression levels (RPKM) of TEs were calculated using reads that were mapped to the TE consensus sequence. Expression levels (RPKM) of coding genes were calculated using TopHat 2.0.14 (Trapnell et al., 2009) and DESeq2 (Love et al., 2014). *Drosophila melanogaster* (dm3) GTF files obtained from Illumina iGenomes were used to define known gene structures. Analysis of coding genes was performed using duplicate samples. Duplicate data were obtained for EGFP-, Nxf2-, and Panx-KD samples, and previously published data (Iwasaki et al., 2016) were used as duplicate data for Piwi-KD samples.

**ChIP-seq analysis**

ChIP was performed as described previously (Iwasaki et al., 2016). Briefly, OSCs ($3\times10^7$) were fixed, lysed, and their nuclei were isolated using truChIP Chromatin Shearing Kits (Covaris), in accordance with the manufacturer's instructions. Sodium deoxycholate was added to a final concentration of 0.4% prior to shearing. DNA was sonicated to ~300 bases using Bioruptor (Cosmobio), and then diluted with ChIP buffer [1× 50 mM HEPES-KOH pH 7.4, 150 mM NaCl, 1 mM EDTA, 1% Triton X-100, Halt Protease Inhibitor Cocktail (Thermo Scientific)]. DNA–protein complexes were incubated with 3 μg of anti-H3K9me3 antibody (61013; Active Motif) on Dynabeads-Protein G for 4 h, at 4°C. Beads were washed with ChIP buffer, high-salt lysis buffer (50 mM HEPES-KOH pH 7.4, 450 mM NaCl, 1 mM EDTA, 1% Triton X-100, 0.1% SDS), and wash buffer (50 mM Tris-HCl pH 8.0, 1 mM EDTA, 250 mM lithium chloride, 0.5% NP-40, 0.5% SDS) followed by TE (10 mM Tris-HCl pH 8.0, 1 mM EDTA). After reversing cross-linking for 12–16 h at 65°C, samples were treated with RNase for 30 min at 37°C and Proteinase K for 60 min at 55°C. DNA was then recovered. To prepare ChIP-seq libraries, DNA fragments from the ChIP experiment were sheared to ~200 bases using Covaris S220 (Covaris). These were used for library preparation with the NEBNext Ultra DNA Library Prep Kit for Illumina II (NEB), following the manufacturer's protocol. ChIP-seq libraries were sequenced with Illumina MiSeq using MiSeq Reagent Kit v3 for 150 cycles (Illumina). For ChIP-seq analysis, adapters were removed from the reads using Cutadapt, and subsequent reads shorter than 50 nt were discarded. ChIP-seq reads were mapped to the *Drosophila* genome (dm3) using Bowtie 1.0.1 (Langmead et al., 2009), using -v 2 --best parameters. Reads mapped to the genome were extracted and mapped to *Drosophila melanogaster* TE consensus sequences from Repbase (Jurka, 1998), permitting only those that mapped uniquely to the TE consensus sequence. Metaplots of ChIP signals were calculated using ngs.plot (Shen et al., 2014).

Coordinates of TE insertions, and the euchromatin/heterochromatin distribution within OSCs were obtained from a previous publication (Sienski et al., 2012). For peak calling of H3K9me3 signals, MACS-1.4.2 (Zhang et al., 2008) was used with the default parameters. ChIP-seq analysis was performed in duplicates for Panx and Nxf2 4-day knockdown samples, and EGFP (control) and Piwi 4-day knockdown samples were obtained from a previous study (Iwasaki et al., 2016).

**CLIP-seq**

For the preparation of preadenylated 3′ adapter, DNA oligonucleotide (5′-/5phos/TGGAATTCTCGGGTGCCAAGG/3Bio/-3′; /5′phos/ = 5′ phosphorylation, /3′Bio/ = 3′ biotin) was custom-synthesized by IDT, and then preadenylated using the 5′ DNA Adenylation Kit (NEB) following the manufacturer's protocol. The reaction was ethanol-precipitated and dissolved in water to a final concentration of 20 μM. cDNA synthesis primer (5′-/5phos/NNAACCNNNGATCGTCGGACTGTAGAACTCTGAA/iSp18/CACTCA/iSp18/ATCTCCT TGGCACCCGAGAATTCCA-3′; /iSp18/ = Carbon-18 spacer) and Illumina sequencing primers [RP1: AATGATACGGCGACCACCGAGATCTACACGTTCAGAGTTCTACAGTCCGA, RPIx: CAAGCAGAAGACGGCATACGAGATXXXXXXGTGACTGGAGTTCCTTGGCACCCGAGAATTCC A, 6-bases index sequence in bold (see Illumina customer service letter for other index sequences)] were custom-synthesized by IDT.

The iCLIP libraries were prepared as previously described (Flynn et al., 2015; Van Nostrand et al., 2016) with the following modifications. For iCLIP from stable cell line of myc-Nxf2, three confluent 15 cm plates (Nunc) per replicate were used. Cells were UV-crosslinked with irradiance of 200 mJ/cm$^2$ at 254 nm using UV Stratalinker 1800 (Stratagene). Cells were harvested and lysed with 2 mL of lysis buffer [20 mM HEPES-KOH (pH7.3), 150 mM NaCl, 1 mM EDTA, 1 mM DTT, 0.5% NP-40, 2 μg/ml pepstatin, 2 μg/ml leupeptin, 0.5% aprotinin]. Cell extracts were treated with 1 U/μL RNase T1 (Roche) for 15 min at room temperature. The lysate was cleared by centrifugation at 13,500 rpm for 15 min at 4°C. The resulting supernatant was then collected and cooled on ice. 10 μL was reserved for size-matched input (SMInput) (Van Nostrand et al., 2016). RNA–protein complexes were immunoprecipitated using 40 μg of mouse monoclonal antibodies (anti-myc 9E10, DSHB) and 200 μL of Protein G Dynabeads (Invitrogen) for 2 h, at 4°C. Beads were then washed three times with IP wash buffer [20 mM HEPES-KOH, (pH7.3), 300 mM NaCl, 1 mM DTT, 0.05% NP-40, 2 μg/mL pepstatin, 2 μg/mL leupeptin, 0.5% aprotinin] and three times with high-salt wash buffer [20 mM HEPES-KOH (pH7.3), 500 mM NaCl, 1 mM DTT, 0.05% NP-40, 2 μg/mL pepstatin, 2 μg/mL leupeptin, 0.5% aprotinin]. Washed beads were resuspended in 30 μL of 1×NuPAGE LDS sample buffer (Thermo Fisher) with 50 mM DTT and incubated at 70°C for 10 min. Released RNA-protein complexes were separated on the 4-12% NuPAGE Bis-Tris gel (Thermo Fisher) with MOPS running buffer (Thermo Fisher) at 200 V for 1 h. After the run, RNA–protein complexes from the gel were transferred to a nitrocellulose membrane using the Novex wet transfer apparatus following the manufacturer's protocol (Thermo Fisher). The transfer was performed for 1 h at 30 V using NuPAGE transfer buffer (Thermo Fisher). Western blotting was performed using SNAP i.d. 2.0 System (Merck) following the manufacturer's instruction to visualize bands and determine regions for isolating RNA–protein complexes. Briefly, the membrane was blocked with 0.5% skim milk in PBS-0.1% Tween 20, incubated with 1:1000 anti-myc antibody, washed with PBS-0.1% Tween 20, and incubated with 1:5000 HRP-conjugated secondary antibody (Cappel). Membranes were developed with ECL substrate (GE) and exposed using film. A region 50 kDa above the protein size was cut into 1–2-mm-narrow strips and then transferred to a 1.5-mL tube. To each tube, 200 μL of Proteinase K buffer [100 mM Tris-HCl (pH 7.5), 150 mM NaCl, 12.5 mM EDTA, 2% (w/v) SDS] and 10 μL of Proteinase K (Roche) was added and incubated for 30 min at 55°C. The RNA was recovered by acidic phenol/chloroform extraction followed by a chloroform extraction and an ethanol precipitation with Pellet Paint NF Co-Precipitant (Merck Millipore). Finally, the pellet was dissolved in 16 µl of nuclease-free water. The recovered RNA was mixed with dephosphorylation mix [2 µL of 10×PNK buffer pH 6.0 (500 mM imidazole-HCl pH 6.0, 100 mM $MgCl_2$, 50 mM DTT), 1 µL of RNasin (Promega), 1 µL of T4 PNK (NEB)] and incubated for 30 min at 37°C. The sample was incubated with 10 µL of SPRIselect (Beckman Coulter) and 30 µL of isopropanol for 5 min and then separated on the magnetic stand for 5 min. The supernatant was discarded and the beads were washed twice with 500 µL of 85% ethanol. The beads were then air dried for 5 min and eluted with 10.2 µL of nuclease-free water. The recovered RNA was mixed with 3′ adapter ligation mix (1 µL of 20 µM preadenylated 3′ adapter, 2 µL of 10×T4 RNA ligase buffer, 4.8 µL of 50% PEG8000, 1 µL of RNasin, 1 µL of T4 RNA ligase2, truncated KQ (NEB)) and incubated overnight at 16°C. Ligation products were purified using 10 µL of SPRIselect and 10 µL of isopropanol in the same procedure as above, and eluted with 16 µL of nuclease-free water. The sample was mixed with 5′ deadenylation mix (2 µL of 10×NEB2 buffer, 1 µL of RNasin, 1 µL of 5′ Deadenylase (NEB)) and incubated 30 min at 30°C. To digest unligated adapters, 5 µL of $RecJ_f$ exonuclease (NEB) was added and incubated 30 min at 37°C. Ligation products were purified using 12.5 µL of SPRIselect and 12.5 µL of isopropanol in the same procedure as above, and eluted with 12.2 µL of nuclease-free water. 1 µL of 10 µM cDNA synthesis primer was added to the whole isolated RNA and annealed to the adapter by incubation at 70°C for 5 min and at 25°C for 1 min. RT mix [4 µL of 5×Superscript III RT buffer, 0.8 µL of dNTP mix (10 mM each), 1 µL of 0.1 M DTT, 1 µL of Superscript III RT (Thermo Fisher)] was added and a reverse transcription reaction was performed using the following program: 42°C for 40 min, 50°C for 15 min, and 4°C for hold. After reverse transcription reaction, 0.5 µL of RNase (Roche) and 0.5 µL of RNaseH (NEB) was added to each sample and incubated at 37°C for 15 min. During this time 5 µL of Dynabeads M-270 Streptavidin (Invitrogen) were washed twice with 200 µL of StrepBead wash buffer (100 mM Tris–HCl, pH 7, 1 M NaCl, 10 mM EDTA, 0.1% Tween-20), for each sample. Each 5 µL volume of beads were finally resuspended in 40 µL of StrepBead wash buffer. After RNase digestion, 40 µL of pre-washed beads were added to each sample and incubated at 25°C for 30 min with rotation. After incubating, the samples were applied to a magnetic stand and the buffer was discarded. Each sample was washed four times with 100 µL of wash buffer (100 mM Tris–HCl pH 7, 4 M NaCl, 10 mM EDTA, 0.2% Tween-20) and twice with 100 µL of NT2 buffer (50 mM Tris–HCl, pH 7.5, 150 mM NaCl, 1 mM $MgCl_2$, 0.05% NP- 40). The purified cDNA was then circularized by adding 20 µL of CircLigase Reaction Mix (2 µL of 10×CircLigaseII Reaction Buffer, 1 µL of 50 mM $MnCl_2$, 1 µL of CircLigaseII ssDNA Ligase (Epicentre), 16 µL of nuclease-free water) to each sample, and incubating for 1 h at 60°C, 10 min at 80°C, and hold at 4°C. Samples were then applied to a magnetic stand, the reaction buffer was collected (be sure not to discard the reaction buffer), and another 20 µL of elution buffer (10 mM Tris pH 7.5 and 1 µM PRIx primer) was added to the beads and heated at 95°C for 2 min. Samples were then applied to a magnetic stand and the elution buffer was removed and combined with reaction buffer, collected earlier (20 µL). The elution was repeated for two times in total obtaining a final volume of 60 µL. cDNA was purified using 30 µL of SPRIselect and 60 µL of isopropanol in the same procedure as above, and eluted with 29.5 µL of nuclease-free water. PCR mix [2.5 µL of 10 µM RP1 primer, 2.5 µL of 10 µM RPIx primer, 10 µL of 5×Q5 Reaction Buffer, 5 µL of dNTP mix (2 mM each), 0.5 µL of Q5 DNA Polymerase (NEB)] was added to the cDNA and PCR was performed using the following program: 98°C for 30 s; cycles of 98°C for 10 s, 60°C for 30 s, and 72°C for 15 s; followed by 4°C hold). DNA was amplified with 10 and 12 cycles of PCR for SMInput samples and immunoprecipitated samples, respectively. Amplified DNA was captured with 1.8 volumes of AMPure XP beads (Beckman Coulter) following the manufacturer's protocol and size-selected on 6% PAGE gels. DNA was visualized with SybrGold (Thermo Fisher) and those with sizes between 170 and 200 bp were isolated. The PAGE gel was crushed, and the DNA was eluted in 400 μL of 0.4 M NaCl at 4°C on rotation overnight. The eluted DNA was ethanol-precipitated with Pellet Paint NF Co-Precipitant and then re-amplified with the same PCR program and for 8 cycles each. PCR reactions were captured with 1.8 volumes of AMPure XP beads, size-selected on 6% PAGE gels, and purified as described above. 1 μL of libraries were quantitated by HS-DNA Bioanalyzer. Libraries were sequenced on the Illumina Miseq platform.

For Piwi iCLIP from OSCs, one confluent 10 cm plate per replicate was used. Cells were lysed in 1 mL of lysis buffer and immunoprecipitation was performed using 10 μg of anti-Piwi antibody and 50 μL of Protein G Dynabeads. The rest of the protocol was identical to that of myc-Nxf2.

**Isolation of myc-Nxf2 stable line OSCs**

Myc-Nxf2 OSC line was isolated using the methodology described in "Tethering assay using OSCs" section, for isolation of OSC line carrying reporter gene. Briefly, OSCs were co-transfected with pAcM-Nxf2-sires plasmid and the plasmid carrying a blasticidin-resistance gene, and selection was performed by blasticidin.

**Shotgun proteome data analysis**

Shotgun proteome was performed in two biological replicates, and two technical replicates per each sample (in total of four measurements per sample). The obtained peptide numbers for four measurements were used to calculate Enrichment and Specificity of obtained protein candidate. Enrichment was calculated by dividing the total number of peptides obtained in Flag-Nxf2-IP samples by control samples, and taking log2. Specificity was calculated as p-value obtained from t-test of peptide counts obtained for Flag-Nxf2-IP samples against control-IP samples.

**Small RNA-seq analysis**

Small RNA cloning was performed as previously described (Iwasaki et al., 2017), with modifications. In summary, coimmunoprecipitated piRNA was extracted from Piwi immunoprecipitates using phenol and chloroform, and purified piRNAs were cloned using the NEXTflex Small RNA Sequencing kit v3 (NEXTflex). The libraries were analysed on a Miseq platform (Illumina).

For small RNA-seq data analysis, adapter sequences were removed from the obtained reads using Cutadapt, and reads out of 20-35nt size range were removed for further analysis. Obtained reads were mapped to the Release 3 assembly of the *Drosophila* genome using Bowtie (Langmead et al., 2009), allowing up to single mismatch. Genome-mapped reads were further mapped to TE consensus sequences from Repbase (Jurka, 1998) using Bowtie (Langmead et al., 2009), allowing no mismatch and unique mapping. The reads were normalized to RPM by the number of *Drosophila* genome-mapped reads. Reads mapped to antisense direction of TE consensus sequences were used to generate heatmap in Figure 6E, and reads mapped to both directions were used to generate density plot in Figure 6F. Sense reads were plotted in positive direction, where antisense reads were plotted in negative direction.

**CLIP data analysis**

First, we trimmed 3′ adapter sequences using Cutadapt and discarded reads shorter than 27 bp. Next, we removed PCR duplicates by collapsing all identical reads containing the same random barcode at the 5′ end with fastx-collapser from the FASTX-Toolkit (http://hannonlab.cshl.edu/fastx_toolkit/index.html). Finally, we trimmed the 5′-end random barcode located at positions 1–9 of the reads. Mapping was then performed against the Release 3 assembly of the *Drosophila melanogaster* genome with STAR (Dobin et al., 2013). Alignment output BAM files were sorted and indexed by SAMtools (Li et al., 2009). Peaks were called using Piranha peak-calling algorithm (Uren et al., 2012), and reads within the peaks were counted for two replicate samples. Of two replicate samples, peak position of the sample that had higher number of annotated peaks was used for counting reads. Dot plots were generated based on the read counts, and correlation coefficient (*r*) was calculated to check for the reproducibility (shown in Figure S6E). After checking for reproducibility, samples were merged for downstream analysis using SAMtools (Li et al., 2009). Genome-mapped reads were further mapped to TE consensus sequences from Repbase (Jurka, 1998) using Bowtie allowing only unique mapping reads. Mapped reads were counted for each consensus TE, and normalized to RPM by the number of *Drosophila* genome-mapped reads. Reads mapped to sense and antisense direction of TE consensus sequences were counted separately, when calculating RPM. Based on the results shown in Figures 6A-C and Figure S6F, sense-mapped read counts were used for analysis shown in Figure 6D-F. Note that piRNAs were not included in the reads mapped to antisense direction of TEs, since the libraries were generated using RNA in the size range of 43~73nt (~50kDa above the protein size). In case of taking ratio of CLIP tags and SMInput, TEs that had RPM lower than 0.5 in SMInput was eliminated. Maximum x-axis for the density plot of Piwi CLIP tags in *mdg1* (Figure 6F) was adjusted to 100, since there was single high peak at position 6870bp (~600 RPM), which masks the other peaks. Dot plot, Box plot, and Density plot was generated using the methodology described in "RNA-seq" and "ChIP-seq" section. Heatmaps were depicted using Java Treeview (Saldanha, 2004).

Saito, K., Inagaki, S., Mituyama, T., Kawamura, Y., Ono, Y., Sakota, E., Kotani, H., Asai, K., Siomi, H.,

and Siomi, M.C. (2009). A regulatory circuit for piwi by the large Maf gene traffic jam in Drosophila. Nature *461*, 1296-1299.
Saito, K., Ishizu, H., Komai, M., Kotani, H., Kawamura, Y., Nishida, K.M., Siomi, H., and Siomi, M.C. (2010). Roles for the Yb body components Armitage and Yb in primary piRNA biogenesis in Drosophila. Genes & development *24*, 2493-2498.
Saito, K., Nishida, K.M., Mori, T., Kawamura, Y., Miyoshi, K., Nagami, T., Siomi, H., and Siomi, M.C. (2006). Specific association of Piwi with rasiRNAs derived from retrotransposon and heterochromatic regions in the Drosophila genome. Genes & development *20*, 2214-2222.
Saldanha, A.J. (2004). Java Treeview--extensible visualization of microarray data. Bioinformatics (Oxford, England) *20*, 3246-3248.
Sato, K., Nishida, K.M., Shibuya, A., Siomi, M.C., and Siomi, H. (2011). Maelstrom coordinates microtubule organization during Drosophila oogenesis through interaction with components of the MTOC. Genes & development *25*, 2361-2373.
Shen, L., Shao, N., Liu, X., and Nestler, E. (2014). ngs.plot: Quick mining and visualization of next-generation sequencing data by integrating genomic databases. BMC genomics *15*, 284.
Sienski, G., Batki, J., Senti, K.A., Donertas, D., Tirian, L., Meixner, K., and Brennecke, J. (2015). Silencio/CG9754 connects the Piwi-piRNA complex to the cellular heterochromatin machinery. Genes & development *29*, 2258-2271.
Sienski, G., Donertas, D., and Brennecke, J. (2012). Transcriptional silencing of transposons by Piwi and maelstrom and its impact on chromatin state and gene expression. Cell *151*, 964-980.
Sumiyoshi, T., Sato, K., Yamamoto, H., Iwasaki, Y.W., Siomi, H., and Siomi, M.C. (2016). Loss of l(3)mbt leads to acquisition of the ping-pong cycle in Drosophila ovarian somatic cells. Genes & development *30*, 1617-1622.
Trapnell, C., Pachter, L., and Salzberg, S.L. (2009). TopHat: discovering splice junctions with RNA-Seq. Bioinformatics (Oxford, England) *25*, 1105-1111.
Uren, P.J., Bahrami-Samani, E., Burns, S.C., Qiao, M., Karginov, F.V., Hodges, E., Hannon, G.J., Sanford, J.R., Penalva, L.O., and Smith, A.D. (2012). Site identification in high-throughput RNA-protein interaction data. Bioinformatics (Oxford, England) *28*, 3013-3020.
Van Nostrand, E.L., Pratt, G.A., Shishkin, A.A., Gelboin-Burkhart, C., Fang, M.Y., Sundararaman, B., Blue, S.M., Nguyen, T.B., Surka, C., Elkins, K.*, et al.* (2016). Robust transcriptome-wide discovery of RNA-binding protein binding sites with enhanced CLIP (eCLIP). Nature methods *13*, 508-514.
Vourekas, A., Zheng, K., Fu, Q., Maragkakis, M., Alexiou, P., Ma, J., Pillai, R.S., Mourelatos, Z., and Wang, P.J. (2015). The RNA helicase MOV10L1 binds piRNA precursors to initiate piRNA processing. Genes & development *29*, 617-629.
Yashiro, R., Murota, Y., Nishida, K.M., Yamashiro, H., Fujii, K., Ogai, A., Yamanaka, S., Negishi, L., Siomi, H., and Siomi, M.C. (2018). Piwi Nuclear Localization and Its Regulatory Mechanism in Drosophila Ovarian Somatic Cells. Cell reports *23*, 3647-3657.
Yu, Y., Gu, J., Jin, Y., Luo, Y., Preall, J.B., Ma, J., Czech, B., and Hannon, G.J. (2015). Panoramix enforces piRNA-dependent cotranscriptional silencing. Science (New York, NY) *350*, 339-342.
Zhang, Y., Liu, T., Meyer, C.A., Eeckhoute, J., Johnson, D.S., Bernstein, B.E., Nusbaum, C., Myers, R.M., Brown, M., Li, W.*, et al.* (2008). Model-based analysis of ChIP-Seq (MACS). Genome biology *9*, R137.
